# Dual polarity voltage imaging reveals subthreshold dynamics and concurrent spiking patterns of multiple neuron-types

**DOI:** 10.1101/2021.10.13.463730

**Authors:** Madhuvanthi Kannan, Ganesh Vasan, Simon Haziza, Cheng Huang, Radek Chrapkiewicz, Junjie Luo, Jessica A. Cardin, Mark J. Schnitzer, Vincent A. Pieribone

## Abstract

Genetically encoded fluorescent voltage indicators are ideally suited to reveal the millisecond-scale interactions among and between distinct, targeted cell populations. However, current indicator families lack the requisite sensitivity for *in vivo* multipopulation imaging. We describe high-performance green and red sensors, Ace-mNeon2 and VARNAM2, and their reverse response-polarity variants, pAce and pAceR. Our indicators enable 0.4-1 kHz voltage recordings from >50 neurons per field-of-view in awake mice and ∼30-min continuous imaging in flies. Using dual-polarity multiplexed imaging, we uncovered behavioral state-dependent interactions between distinct neocortical subclasses, as well as contributions to hippocampal field potentials from non-overlapping projection neuronal ensembles. By combining three mutually compatible indicators, we demonstrate concurrent triple-population voltage imaging. Our approach will empower investigations of the dynamic interplay between neuronal subclasses at single-spike resolution.

**One Sentence Summary:** A new suite of voltage sensors enables simultaneous cellular-resolution activity imaging from multiple, targeted neuron-types in awake animals.

## Main

The dynamic interplay of multiple excitatory and inhibitory neuronal subclasses lies at the core of how the brain processes information. In context-dependent motor and sensory processing in neocortex, the external environment or the animal’s internal behavioral state impacts the activity of local interneurons, which in turn modulates the outputs of excitatory cells (*1-5*). Likewise, in the hippocampus, distinct projection neuronal ensembles exhibit state-dependent changes in firing rates to differentially encode learning and memory (*6-9*). However, how different subclasses coordinate their activity patterns in real-time to control network output remains elusive due to an inability to simultaneously measure the voltage dynamics within and across targeted ensembles.

Genetically encoded voltage indicators (GEVIs) enable targeted recordings of spiking and subthreshold activity with sub-millisecond temporal precision, offering unique advantages over microelectrodes and Ca^2+^ imaging (*10-12*). Although extensive protein engineering has uncovered a host of new sensors (*13-21*), existing GEVIs fall short of enabling recordings from large neural ensembles in behaving animals. Notably, *Archaerhodopsin*-based sensors are extremely dim and require high illumination powers of >1 W mm^-2^ under widefield microscopy or, alternatively, need specialized apparatus providing patterned illumination to improve signal-to-noise ratios and to record from tens of cells simultaneously (*16, 19, 22-24*). Fluorescence Resonance Energy Transfer (FRET)-opsin indicators, which combine opsins with bright genetic or dye-based fluorescent proteins (FPs), are more easily scalable to large populations but exhibit modest sensitivities or require repeated administrations of exogenous chemical dyes (*13, 14, 17, 18*).

We previously introduced the green and red FRET-opsin GEVIs Ace-mNeon and VARNAM, which are molecular fusions of the voltage-sensitive *Acetabularia* (Ace) rhodopsin with the bright FPs mNeonGreen and mRuby3, respectively (*17, 18*). FRET-opsins that are fully genetically encoded overcome the constraints of other GEVI families, and, notably, Ace-mNeon and VARNAM remain to date the only pair of mutually compatible sensors allowing concurrent recordings in distinct cell classes (*18*). Here, to enable large-scale simultaneous multipopulation imaging, we engineered next-generation GEVIs, Ace-mNeon2 and VARNAM2, and their corresponding reverse response-polarity variants, pAce and pAceR, all of which are mutually compatible for *in vivo* voltage imaging.

Our improved indicators have >50% enhanced sensitivities over their predecessors and enable low-power (<25 mW mm^-2^) recordings from >50 neurons per field-of-view in awake mice and up to 30-min continuous imaging in flies. We further combine the normal and reverse polarity indicators for dual-polarity multiplexed voltage imaging (DUPLEX) to uncover the millisecond-scale interactions *between* cortical subtypes during behavioral state transitions. We also unveiled real-time relationships between two concurrently recorded hippocampal subtypes and the local field potential (LFP). Lastly, by combining DUPLEX with dual color imaging, we achieve simultaneous cellular-resolution activity recordings from as many as 57 neurons belonging to three distinct subclasses, providing the first insights into subclass-specific voltage dynamics in intact networks.

### Protein engineering of negative and positive polarity FRET-opsins

To improve the sensitivities of Ace-mNeon and VARNAM for live animal imaging, we performed protein engineering combined with high-throughput voltage screening in excitable human embryonic kidney (HEK) cells (Fig. 1,A and B) (*18, 25, 26*). Our semi-automated screening platform scored GEVIs for brightness, voltage sensitivity and kinetics in plate-based format, allowing us to screen 120 unique variants/day with a sample content of ∼2000 cells/variant (*18*).

**Fig. 1.**
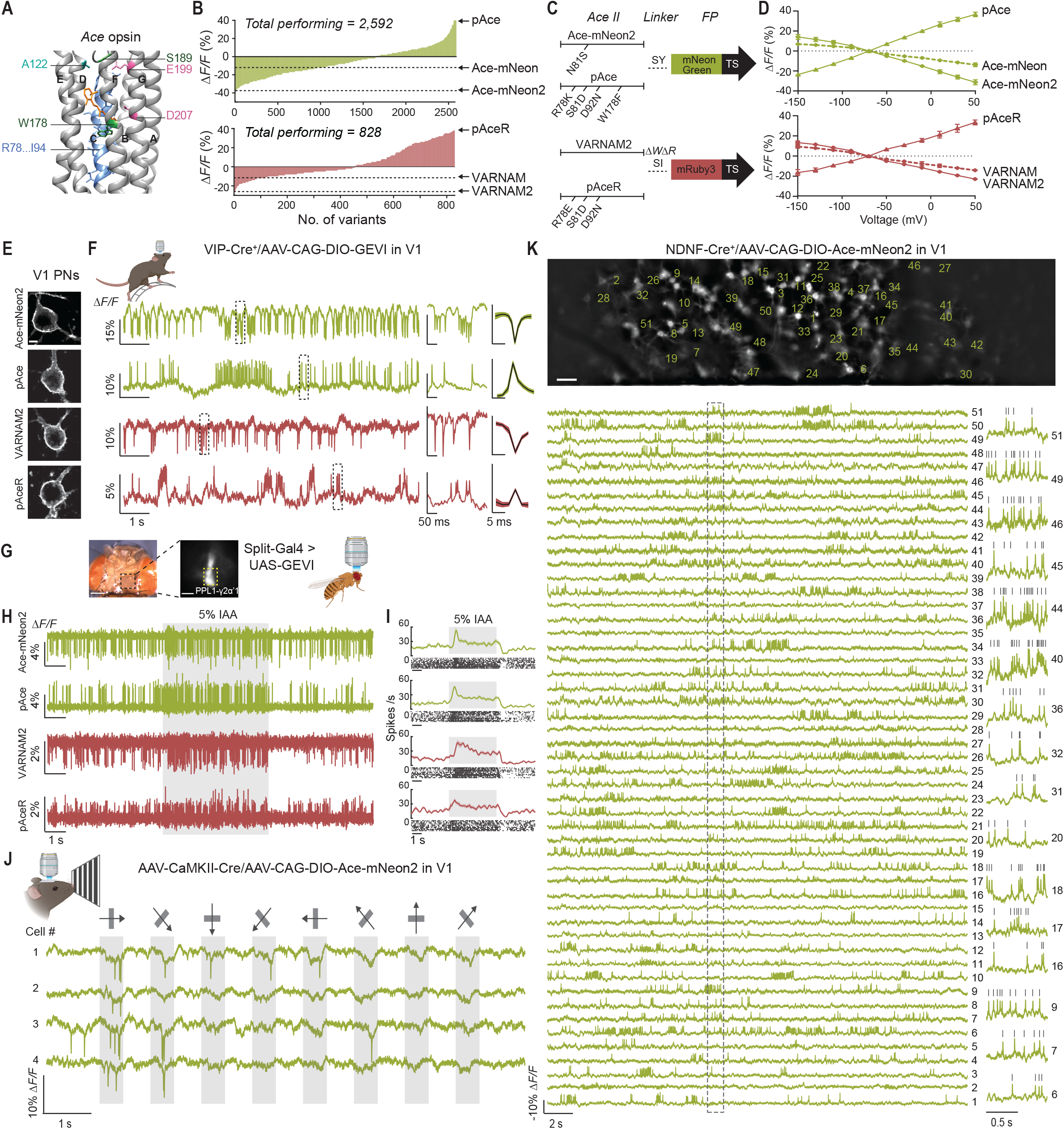
Design and characterization of Ace-mNeon2, VARNAM2 and reverse polarity variants pAce and pAceR. (A) Crystal structure of Ace (PDB ID: 3AM6) showing residues targeted for site-directed saturation mutagenesis on Ace-mNeon and VARNAM. The 7 transmembrane helices are labeled A through G. AAs R78 through I94 on helix C and specified loci on other helices were targeted. (B) Distribution of fluorescence responses to field stimulation acquired from spiking HEK cells expressing Ace-mNeon (*top*) and VARNAM (*bottom*) variants obtained on the high-throughput platform. (C) Schematic representation of Ace-mNeon2, pAce, VARNAM2 and pAceR constructs depicting the point mutations in Ace and Ace-FP linker. (D) *ΔF/ΔV* curves obtained using whole-cell recordings with concurrent fluorescence imaging of HEK cells transfected with Ace-mNeon, Ace-mNeon2 and pAce (*top*) and VARNAM, VARNAM2 and pAceR (*bottom*). Values represent mean ± S.E.M. (E) Confocal images of V1 PNs expressing the indicated sensors. Scale bar: 5 μm. (F) *Left*, example fluorescence traces showing spontaneous spiking obtained from awake, head-restrained mice selectively expressing the indicators in V1 VIP-interneurons; *Center*, the boxed areas on the traces are shown at an expanded timescale; *Right*, Mean ± S.E.M. optical spike waveform. (G) *In vivo* voltage imaging through a transparent surgical window implanted on a fly head (*left*). Each of the indicators was selectively expressed in PPL1-γγ2α’1 dopaminergic neuron using a split-GAL4 system (*right*). Yellow dashed box indicates the axonal region of PPL1-γγ2α’1. (H) Example fluorescence recordings obtained using the four indicators showing odor-evoked spiking elicited by 5 s delivery of 5% isoamyl acetate in PPL1-γγ2α’1. (I) Mean ± S.E.M. firing rate change during odor delivery and raster plots of individual trials (n = 10 trials, 2 trials/fly). (J) Example visual responses from layer 2/3 PNs to drifting gratings presented at eight different orientations acquired using Ace-mNeon2 in an awake mouse. Arrows indicate stimulus orientation and grey shaded regions correspond to stimulus periods. (K) *Top*, Representative epifluorescence image of a single field-of-view from a NDNF-*Cre*^*+*^ mouse expressing *Cre*-dependent soma-targeted Ace-mNeon2 in V1. Scale bar: 50 μm. *Bottom, ΔF/F* traces showing spontaneous activity from regions-of-interest (ROIs) numbered in the image above. Fluorescence traces are inverted for visualization. Boxed region is shown at an expanded timescale on the right for select cells. Grey ticks denote identified spikes.

In Ace-mNeon and VARNAM, the absorption spectrum of Ace (FRET acceptor) overlaps with the emission spectrum of the FP (FRET donor), allowing for depolarization-dependent fluorescence quenching by the opsin, which is seen as fluorescence *decreases* in the optical readout. To improve the voltage sensitivities of the indicators, we therefore sought to enhance FRET transfer by optimizing the Ace-FP linker (*27*). In Ace-mNeon, we further performed site-saturation mutagenesis at Ace N81, owing to its contribution to VARNAM’s voltage sensitivity (Fig. 1A, and S1) (*18*). Screening of mutagenic libraries uncovered variants with increased response amplitudes (Fig. 1B, and S2). Ace-mNeon N81S G229Y and VARNAM *Δ*228W *Δ*229R G231I (henceforth, Ace-mNeon2 and VARNAM2, respectively) exhibited significant improvements in fluorescence responses to depolarizing voltages compared to their predecessors in patch clamp recordings from transfected cells (*ΔF/F* for 120 mV step were (mean ± S.E.M.): − 31.297 ± 0.7% for Ace-mNeon2 (n=13 cells) versus −13.85 ± 0.6% for Ace-mNeon (n=11 cells), *P*<0.001, two-tailed t-test; −22.87 ± 0.7% for VARNAM2 (n=13 cells) versus −14.1 ± 0.5% for VARNAM (n=8 cells), *P*<0.001, two-tailed t-test) (Fig. 1, C, and D, and S3, S4, and S5 and Supplementary table 1).

We next screened a library of Ace-mNeon2 D92X positional mutants, owing to the highly conserved role of the N81/D92 pair in opsin proton-pumping and photosensing (*28, 29*). We uncovered several variants with response polarity reversals, which exhibited fluorescence *increases* during voltage depolarization (Fig. S6). Further, while the top-performing positive variant Ace-mNeon2 D92N exhibited slower kinetics, reciprocal S81X saturation mutagenesis revealed rescue mutants (Fig. S6, and S7). Notably, the 92N/81D reversed polarity phenotype is transposable across other Ace-based indicators, including VARNAM and Voltron (*13, 18*).

For further engineering, we targeted residues lining Ace’s photosensing helix C, amino acids involved in stabilizing its voltage-sensitive intermediate state, and those in physical proximity to the chromophore (Fig. 1A, and S1) (*29-31*). Voltage screening of mutagenic libraries uncovered positive variants with enhanced sensitivities (Fig. S8). Ace-mNeon2 R78K S81D D92N W178F and VARNAM2 R78E S81D D92N, henceforth referred to as pAce (positive Ace) and pAceR (positive Ace in Red), exhibited *ΔF/F* values of 36.6 ± 0.7% (mean ± S.E.M.) and 33.4 ± 1.1% (n=11 cells each), respectively, per 120 mV in cultured HEK cells (Fig. 1, B-D, and S5, S9, and S10). All indicators except pAceR exhibited sub-millisecond kinetics at room temperature (Fig. S5). Additionally, because the positive variants provide the same dynamic range of signaling as Ace-mNeon2 and VARNAM2 but in the opposite direction, they rest at slightly lower intensities than the latter and exhibit reduced resting state photobleaching, akin to other positive GEVIs (Fig. S11) (*14, 32*).

### Characterization of sensors in mice and flies

To characterize our indicators for live animal imaging, we generated soma-targeted versions of Ace-mNeon2, VARNAM2, pAce and pAceR, which trafficked to the neuronal somatic membrane (Fig. 1E) (*33*). All four sensors robustly captured action potentials in vasoactive intestinal peptide (VIP)-expressing interneurons in the primary visual cortex (V1) of head-fixed, awake mice under high-speed (400 Hz) 1-photon fluorescence microscopy (Fig. 1F). The average *in vivo ΔF/F* values per spike were (mean ± S.E.M.): −16.51 ± 1.9% for Ace-mNeon2 (n=127 spikes), 10.67 ± 1.4% for pAce (n=87 spikes), −7.3 ± 1.2% for VARNAM2 (n=49 spikes) and 3.15 ± 0.5% for pAceR, (n=48 spikes).

We further expressed each of the four indicators in PPL1-γ2α’1 dopaminergic neurons in flies, where they reported spontaneous and odor-evoked spiking with comparable baseline- and evoked spike rates (Fig. 1, G-I). The green indicators, compared to the red sensors, exhibited greater response amplitudes and *d’*, a theoretical signal detection metric of spike discriminability (*34*) (*ΔF/F, d’* (mean ± S.E.M.): 3.1 ± 0.2%, 14.54 ± 1.4 (Ace-mNeon2); 4.0 ± 0.1%, 13.4 ± 0.8 (pAce); 1.1 ± 0.1%, 7.4 ± 0.4 (VARNAM2); 0.9 ± 0.03%, 6.5 ± 0.3 (pAceR)). Further, all four indicators showed increased *d’* values compared to parent indicators Ace-mNeon and VARNAM in PPL1-α’2α2 neurons (∼10.8 (Ace-mNeon2), ∼8.7 (pAce), ∼7.0 (VARNAM2) and ∼6.7 (pAceR) versus previously reported values of ∼6.9 (Ace-mNeon) and ∼6.6 (VARNAM)) (Fig. S12) (*18*). Crucially, pAce’s higher *ΔF/F* values and superior photostability enabled the very first extended duration (30-min) continuous recordings at single-spike resolution with consistent *d’* throughout imaging sessions under illumination powers of ∼5 mW mm^-2^ (Fig. S13).

To record stimulus-evoked activity in awake mice, we next expressed our best-performing sensor Ace-mNeon2 in a sparse subset of layer 2/3 pyramidal neurons (PNs) and performed voltage imaging during presentations of drifting grating stimuli. Ace-mNeon2 captured orientation-selective visual responses in fluorescently labeled PNs (Fig. 1J). Further, to demonstrate cellular-resolution large ensemble imaging, we targeted the indicator to cortical layer 1-specific neuron-derived neurotrophic factor (NDNF)-positive interneurons (*35*) and performed widefield voltage imaging. Ace-mNeon2 reported activity from as many as 51 NDNF-neurons in a single field-of-view (Fig. 1K). Likewise, both Ace-mNeon2 and pAce captured voltage signals from multiple cells simultaneously, depending on the labeling density, when selectively expressed in layer 2/3 VIP- and somatostatin (SST)-expressing interneurons (Fig. S14, A, C and E). In these recordings, the *ΔF/F* traces from juxtaposed neurons, <50 μm of each other, were uncorrelated, suggesting minimal cross-contamination of voltage signals (Fig. S14, B, D and F).

### Voltage imaging of visual cortical ensembles during behavioral state transitions

In neocortex, local interneuron activity is strongly influenced by the animal’s internal behavioral state and in turn modulates the output of excitatory cells (*1-5*). However, the extent to which individual subtypes respond to state transitions and how distinct brain states may impact the millisecond-scale interactions between targeted ensembles remain unknown. To study the effect of state on firing rate modulation in distinct cell types, we selectively expressed Ace-mNeon2 in superficial V1 interneurons expressing NDNF, VIP or SST by injecting an adenoassociated virus (AAV) encoding *Cre*-dependent voltage indicator in mouse driver lines expressing *Cre*-recombinase in each of these cell-types (*36, 37*). Additionally, to label a sparse subset of mostly layer 2/3 PNs, we targeted neurons projecting to the anteromedial visual cortical area (AM) (*38, 39*), by delivering the AAV for floxed Ace-mNeon2 in V1 of wild-type mice along with a retrograde CaMKII-*Cre* virus in the AM area. We then performed 400 Hz voltage imaging in V1 of awake, head-restrained mice (Fig. 2A).

**Fig. 2.**
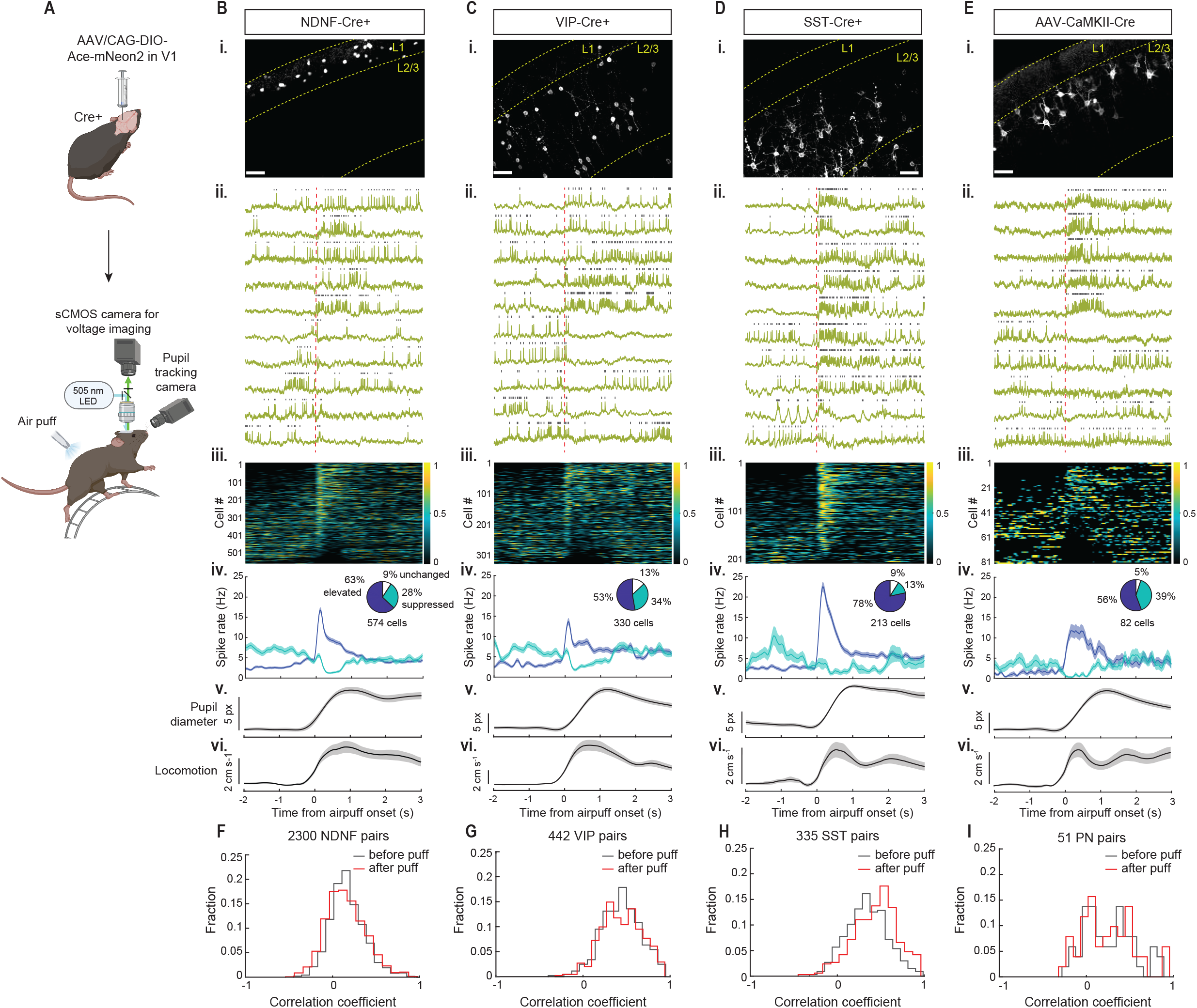
Ace-mNeon2 voltage imaging unveils state-dependent modulation of spontaneous firing in excitatory and inhibitory cell types in V1. (A) Schematic of AAV injection and experimental setup. (B) (i) Confocal images of soma-targeted Ace-mNeon2 expression in NDNF-interneurons. Scale bar: 50 μm. (ii) Representative fluorescence-time traces from individual cells (inverted for visualization purposes). Vertical dashed line in red indicates air puff onset. Grey ticks denote identified spikes. (iii) Z-scored firing rate PSTH for all NDNF-interneurons (n=574 cells, 6 mice). Cells are arranged in order of decreasing spike modulation indices (see Methods). (iv) Mean ± S.E.M. firing rate aligned to air puff onset for activated (dark blue) and suppressed (cyan) fractions. Pie chart insets indicate % of cells with elevated (dark blue), suppressed (cyan) or unchanged (white) spike rates following air puff. (v) Average pupil diameter and (vi) locomotory speed. (C) Same as (B) for VIP-interneurons (n=330 cells, 5 mice). (D) Same as (B) for SST-interneurons (n=213 cells, 6 mice). (E) Same as (B) for PNs (n=82 cells, 4 mice). (F-I) Distribution of pairwise zero time-lag correlation coefficients of the *ΔF/F* traces for (F) NDNF-interneurons (2300 pairs, 6 mice), (G) VIP-interneurons (n=442 pairs, 5 mice), (H) SST-interneurons (n=335 pairs, 6 mice), and (I) PNs (n=51 pairs, 4 mice) computed across a 1 s time window before and after air puff.

The average spontaneous firing rates estimated from the *ΔF/F* traces were (mean ± S.E.M.): 4.72 ± 0.01 Hz for NDNF-interneurons (n=192 cells, 7 mice), 6.52 ± 0.23 Hz for VIP-interneurons (n=49 cells, 5 mice), 4.71 ± 0.21 Hz for SST-interneurons (n=43 cells, 5 mice) and 0.93 ± 0.34 Hz for PNs (n=53 cells, 5 mice). Our values are consistent with the estimated firing rates of V1 layer 2/3 PNs and SST-neurons obtained using electrophysiological recordings in awake mice (*4, 40*). Likewise, the relatively higher spontaneous firing among VIP-compared to SST-interneurons has been observed across distinct cortical areas (*41, 42*).

Next, to study the effect of state during these imaging sessions, we causally induced state change from resting to arousal by delivering a brief air puff to the back of the animal, while simultaneously monitoring the pupil diameter (Fig. 2A) (*43*). Arousal, marked by pupil dilation (*43, 44*), was often accompanied by onset of locomotion (Fig. 2, B-E). A change in cortical state was further confirmed by a decrease in low-frequency subthreshold power and increase in high-frequency subthreshold power, extracted from the *ΔF/F* traces (Fig. S15) (*43, 44*). This led to a robust state-dependent modulation of firing rates across all cell types recorded (Fig. 2, B-E). However, there were important distinctions between cortical subtypes in terms of the relative fraction of cells which increased or decreased their firing, the magnitude of spike rate modulation and response latencies.

Air puff delivery had the most profound effect on SST-interneurons, where the majority (78%) of cells increased and only a small fraction (13%) suppressed their spiking. SST-cells also exhibited the largest increase in spike rate among the four cell types imaged (2.54 ± 0.4 Hz (mean ± S.E.M.) under baseline conditions versus 22.52 ± 1.2 Hz following air puff) (Fig. 2D). They were closely followed by NDNF-interneurons, 63% of which increased their firing rates from an average of 2.56 ± 0.3 Hz under baseline conditions to 16.8 ± 0.7 Hz following air puff, while only 28% decreased their spiking activity (Fig. 2B). Among VIP-interneurons and PNs, comparable fractions of cells elevated (53% and 56%, respectively) or suppressed (34% and 39%, respectively) their firing and both populations showed similar amplitude of spike rate increases during arousal (from 3.75 ± 0.5 Hz to 13.65 ± 0.8 Hz for VIP-cells, and 0.94 ± 0.3 Hz to 11.78 ± 1.6 Hz for PNs) (Fig. 2, C and E). While the elevated firing of SST interneurons fell sharply, as arousal levels peaked and gradually declined, those of VIP- and NDNF-interneurons, which receive strong inhibitory input from SST-cells (*35, 45*), showed an initial rapid decline, followed by a more gradual return to baseline (Fig. 2, B-D).

Next, to estimate the degree of synchronization between neighboring cell pairs, we evaluated zero-time lag correlation coefficients of the *ΔF/F* traces before versus after air puff delivery (Fig. 2, F-I). Both SST- and VIP-interneurons exhibited strong intra-population coupling of spontaneous activity, consistent with prior findings (*46*). These correlations increased slightly during arousal, especially among SST-cells (mean ± S.E.M. correlation coefficients were: 0.37 ± 0.01 under baseline conditions versus 0.44 ± 0.01 following air puff for SST-neurons; 0.43 ± 0.01 versus 0.45 ± 0.01 for VIP-neurons) (Fig. 2, G and H). By comparison, NDNF-interneurons and PNs exhibited weak intra-type correlations (*47*), which were unaltered during state transitions (mean ± S.E.M. correlation coefficients were: 0.17 ± 0.004 versus 0.15 ± 0.005 for NDNF-neurons; 0.24 ± 0.04 versus 0.24 ± 0.04 for PNs) (Fig. 2, F and I).

### Dual polarity multiplexed voltage imaging

While behavioral state differentially modulates the firing rates of distinct cell types (Fig. 2), it remains unknown whether, as a result, state also affects the synergistic activity patterns between two or more distinct subtypes. To examine this question, we turned to simultaneous multipopulation voltage imaging using our improved indicators.

In the past, multipopulation optical recordings using Ca^2+^ or voltage sensors have been achieved with spectrally orthogonal sensors targeted to distinct cell types (*18, 48-51*). However, we reasoned that a pair of GEVIs with opposite response-polarity but operating in the same spectral channel should be mutually compatible, enabling studies of two cell-types per color channel. The cells of each class can be distinguished via the directionality of their optical spike waveforms.

To validate this approach, we performed dual polarity multiplexed voltage imaging (DUPLEX) in awake animals, where we simultaneously targeted Ace-mNeon2 and pAce to two inhibitory cell types, or to one excitatory and one inhibitory cell types, using *Cre* and *Flp*-dependent expression, respectively (Fig. 3A and S16, see also Methods). Selective AAV-mediated expression of a *Cre*-dependent Ace-mNeon2 in NDNF-interneurons and a *Flp*-dependent pAce in V1 VIP-cells in a NDNF-*Cre*^+^/VIP-*Flp*^+^ double transgenic mouse, revealed spontaneous, time-varying spiking and subthreshold dynamics within and between the two cell types (Fig. 3, B and C). In single fields-of-view comprising multiple neurons of both subtypes, VIP-interneurons and, to a lesser extent, NDNF-interneurons exhibited positive intra-population correlation coefficients as noted in Fig. 2, B and C (mean ± S.E.M. correlation coefficients were: 0.31 ± 0.05 for VIP-VIP pairs and 0.14 ± 0.003 for NDNF-NDNF pairs) (Fig. 3, D and E) (*46*). Notably, however, the voltage dynamics were uncorrelated between NDNF-VIP pairs, with a mean inter-type correlation coefficient of −0.09 ± 0.01, suggesting that the two subtypes may receive distinct inputs or that they may not be the principal targets of one another (Fig. 3, D and E) (*35, 52*). Among all pairs, the intercellular correlations decayed as a function of distance between the cells (Fig. S17).

**Fig. 3.**
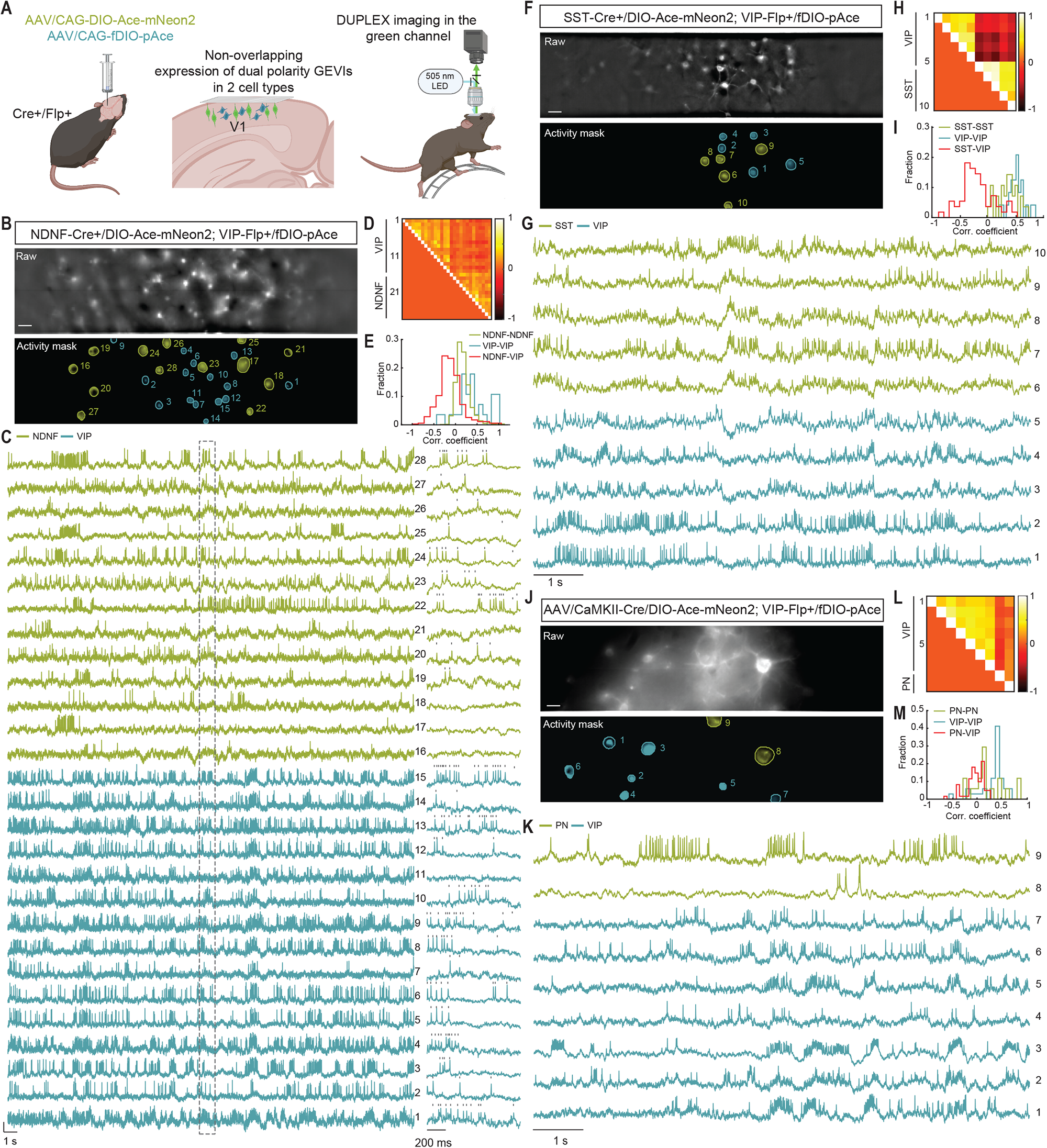
DUPLEX captures the concerted activation dynamics between cell-type pairs in V1. (A) Schematic of AAV injections and experimental setup. (B-M) DUPLEX recordings from two cell populations. (B, F and J) Raw epifluorescence (*top*) and activity-mask (*bottom*) images of single fields-of-view from a (B) NDNF-*Cre*^*+*^*/*VIP-*Flp*^*+*^ mouse expressing Ace-mNeon2 in NDNF-interneurons (green) and pAce in VIP cells (blue), (F) SST-*Cre*^*+*^*/*VIP-*Flp*^*+*^ mouse expressing Ace-mNeon2 in SST interneurons (green) and pAce in VIP-interneurons (blue) and (J) a VIP-*Flp*^*+*^ mouse expressing Ace-mNeon2 in PNs (green) and pAce in VIP-interneurons (blue). Active regions-of-interest (ROIs) are numbered. Scale bar: 50 μm. (C, G and K) *ΔF/F* traces from the ROIs numbered in (B), (F) and (J), respectively. Ace-mNeon2 traces are inverted for visualization purposes. The boxed region in (B) is shown at an expanded timescale to the right. Grey ticks indicate identified spikes. (D, H and L) Intra- and inter-population correlation coefficient matrix constructed from pairwise, zero time-lag correlation coefficients of the *ΔF/F* traces in (C), (G) and (K), respectively. (E, I and M) Distribution of correlation coefficients across all mice for (E) NDNF/VIP recordings (n=717 NDNF-NDNF pairs, 77 VIP-VIP pairs, and 249 NDNF-VIP pairs; 83 NDNF-neurons and 42 VIP-neurons, 5 mice), (I) SST/VIP recordings (n=33 SST-SST pairs, 13 VIP-VIP pairs, and 56 SST-VIP pairs; 17 SST-neurons and 25 VIP-neurons, 3 mice) and (M) PN/VIP recordings (n=17 PN-PN pairs, 34 VIP-VIP pairs, and 61 PN-VIP pairs; 20 PNs and 22 VIP-neurons, 3 mice).

In contrast, in single fields-of-view comprising multiple SST- and VIP-interneurons, intra-population spiking and subthreshold activity were tightly synchronized (mean ± S.E.M. correlation coefficients were: 0.36 ± 0.03 for SST-SST pairs and 0.52 ± 0.03 for VIP-VIP pairs) but the inter-population dynamics were strongly negatively correlated with an average correlation coefficient of −0.24 ± 0.03 among SST-VIP pairs (Fig. 3, F-I). The results are consistent with bidirectional inhibition between the two cell types and the non-overlapping nature of synaptic input to SST and VIP cells (*45, 53*).

Lastly, while we did not observe pairwise zero-time lag inter-population correlations between PNs and VIP-interneurons (mean ± S.E.M. correlation coefficients of 0.31 ± 0.04 for VIP-VIP pairs; ± 0.07 for PN-PN pairs; −0.03 ± 0.02 for PN-VIP pairs) (Fig. 3, J-M), computations of non-zero time-lag cross-correlations revealed a small positive correlation (Fig. S18), likely reflecting the temporal delay in transmission consistent with the lack of direct connections between VIP-neurons and PNs (*45*).

### DUPLEX imaging in subcortical areas

DUPLEX can be extended to study voltage dynamics in subcortical regions such as the hippocampal CA1, where when combined with LFP recordings, it can help unravel the relative contributions of two identified subtypes to the LFP readout (Fig. 4A). In a running mouse expressing Ace-mNeon2 in entorhinal cortex (EC)-projecting excitatory neurons in the *stratum pyramidale* and pAce in SST-interneurons in the *stratum oriens*, we extracted activity from 10 excitatory and 18 SST-cells within the same field-of-view, identifying cell type by the optical spike polarity (see Methods). SST-interneurons exhibited increased baseline firing rate and burstiness compared to excitatory cells (19.8 ± 1.8 Hz versus 14.7 ± 0.5 Hz (mean ± S.E.M.), *P*<0.01) (Fig. 4, B-E), consistent with previous findings. We next assessed the extent to which intrinsic subthreshold or the tissue LFP dynamics organize suprathreshold activity of the excitatory versus inhibitory neurons. Among a total of 157 neurons, we found that spiking from both cell types consistently phase-locked to intra-population subthreshold oscillations compared to tissue LFP: SST-cells and EC-projecting PNs fired at a specific theta (4-9 Hz) phase of their respective transmembrane potentials *V*_*m*_, with distinct average phase tuning of 13.8 ± 2.6° and −1.3 ± 4.8°, respectively (mean ± S.E.M., *P*<0.01) (Fig. 4, F-H). Further, spiking in EC-projecting cells, but not SST-interneurons, consistently phase-locked to the LFP (*P*<0.001 and *P*=0.82, respectively) (Fig. 4, G and H).

**Fig. 4.**
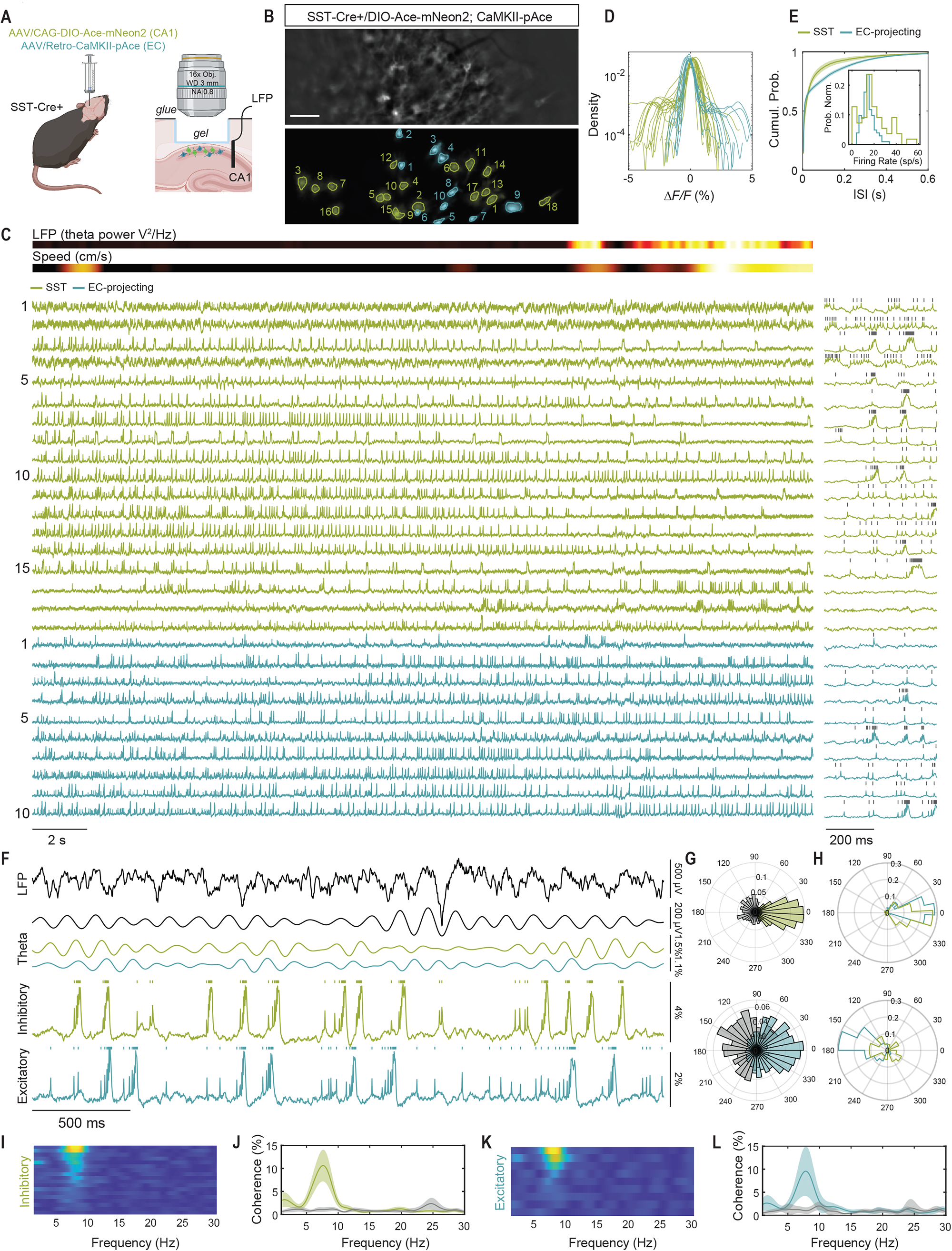
DUPLEX uncovers subtype-specific contributions to LFP in hippocampal CA1. (A) Schematic of AAV injections and experimental setup. (B) Representative raw epifluorescence image (*top*) and spatial footprint of negative- and positive-polarity signals (in green and blue, respectively) of 18 and 10 identified neurons. Scale bar: 50 μm. (C) *ΔF/F* traces for all the neurons in (B) along with wheel speed and LFP theta (*top*). Ace-mNeon2 traces are inverted for visualization purposes. The first 500 ms is shown at the expanded timescale to the right. Grey ticks denote identified spikes. (D) Time course distribution of all neurons across all fields-of-view. Note the opposite polarities within similar dynamic ranges for Ace-mNeon2 and pAce. (E) Cumulative distribution of the inter-spike interval and firing rate (inset) for all recorded neurons (n=55 SST-interneurons and 102 projection neurons, 6 fields-of-view, 1 mouse). Shaded areas represent 95% CI. (F) Phase relationships between spike and theta oscillations extracted from the LFP or cellular transmembrane voltage oscillations, for projection neurons and SST-cells recorded simultaneously. Shown are the raw LFP trace (*top*), theta-filtered (5 Hz-10 Hz) (*center*) and unfiltered excitatory and inhibitory traces (*bottom*). (G) Spike-theta phase relationship for the two neurons in (F). Color and gray represent theta V_m_ and theta LFP, respectively. Note that both neurons are phase-locked to their own V_m_ and LFP but in the opposite phase. (H) Polar histogram of probability density of the average spike-theta phase relationship for all 157 neurons, computed against their respective theta V_m_ (*top*) or theta LFP (*bottom*). (I-L) LFP-subthreshold coherence of all neurons in (C), showing that a fraction of (I and J) SST-interneurons and (K and L) EC-projecting excitatory neurons are phase locked with the LFP at the theta frequency.

We next assessed the cell-type-specific subthreshold contributions to the LFP and the subthreshold relationships within and between two populations. We observed strong subthreshold coherence between both subtypes and LFP in the theta frequency band (*P*<0.0001 and *P*<0.0001, respectively) (Fig. 4, I-L, and S19, A-D). We also observed strong intra- and inter-population coherence in the beta frequency band (15-25 Hz), which was absent in the LFP (Fig. S19, E-H). Overall, these findings are consistent with the prevailing hypothesis that excitatory neurons contribute more strongly to the LFP due to their spatial organization and temporal synchronization (*54*).

### DUPLEX imaging in live flies

To demonstrate the applicability of DUPLEX for multi-cell-type recordings across species, we expressed pAce in GABAergic MBON-γ1pedc>α/β neuron and Ace-mNeon2 in dopaminergic PPL1-α′2α2 in flies (Fig. S20A). While both MBON-γ1pedc>α/β and PPL1-α′2α2 neurons receive excitatory olfactory inputs from mushroom body Kenyon cells, MBON-γ1pedc>α/β itself exerts feedback inhibition on PPL1-α′2α2 axons (*55*). Thus, when we exposed the double-labeled flies to attractive and repulsive odorants, DUPLEX recordings in the two neurons revealed responses of opposite valences in the two neuron-types (Fig. S20, B and C).

### Inter-population correlations of spiking dynamics during state transitions

Having validated DUPLEX for simultaneous dual population recordings, we next applied the technique to uncover the effect of state transitions on the millisecond-scale interactions between distinct subtypes. In DUPLEX recordings from NDNF- and VIP-interneurons, we found that under baseline conditions, spiking and subthreshold activity between the two populations were uncorrelated as seen above (Fig. 3, D and E, and 5, A-D). Arousal only had a small negative impact on inter-population correlations, leading to a slight leftward shift of the distribution of correlation coefficients (mean ± S.E.M. correlation coefficients were: 0.06 ± 0.007 under baseline conditions versus 0.03 ± 0.009 following air puff) (Fig. 5C). We further computed the spike modulation index, defined as (FR_arousal_ − FR_baseline_)/(FR_arousal_ + FR_baseline_), where FR is the firing rate during a 750 ms window before and after air puff delivery, for neighboring pairs of NDNF- and VIP-cells. We found that the paired modulation indices clustered in the top right quadrant, corresponding to a positive modulation for both cell types (Fig. 5D). Our findings suggest a higher propensity for neighboring NDNF- and VIP-cells to increase their firing rates during state transitions.

**Fig. 5.**
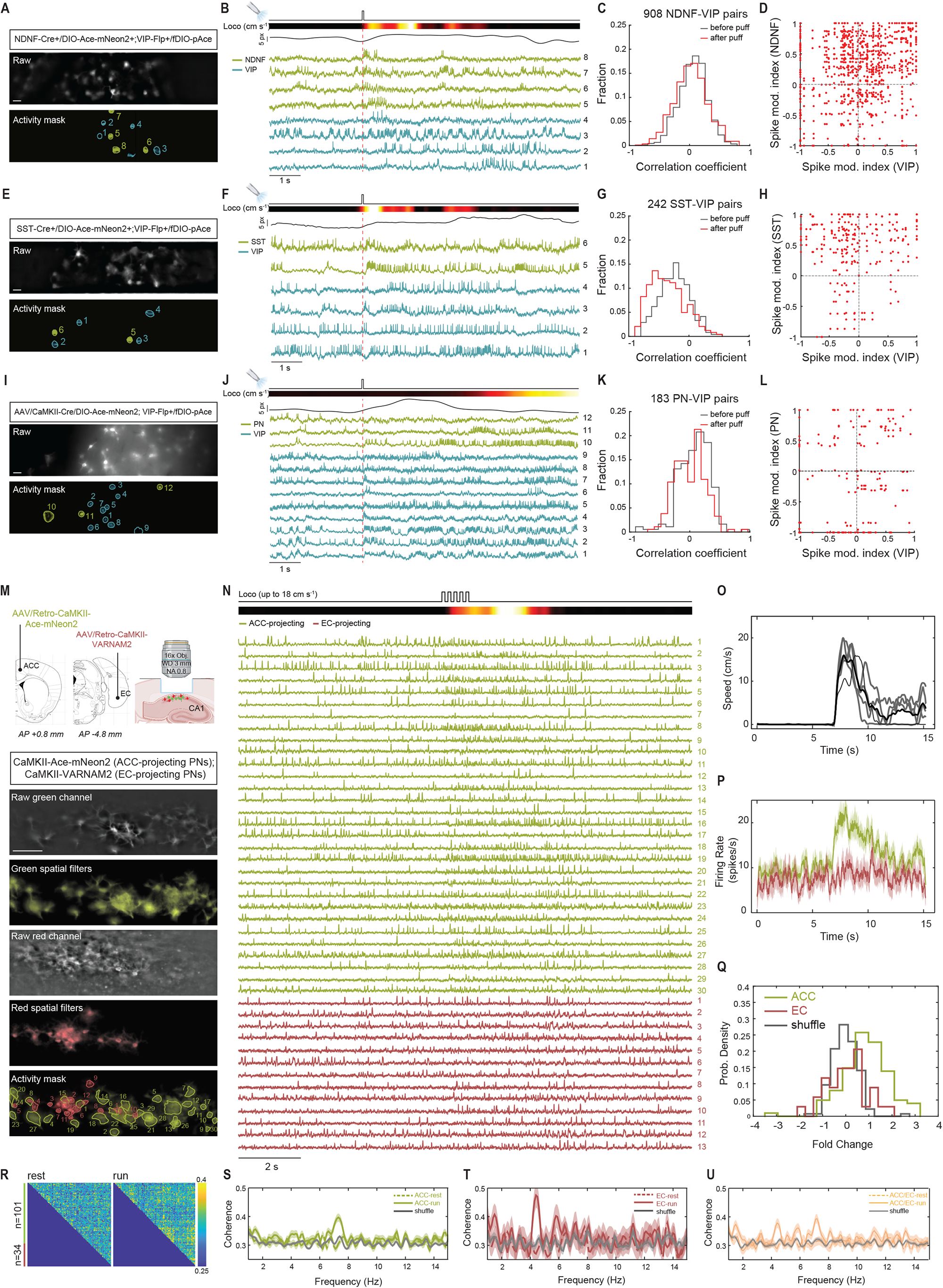
Dual population recordings in V1 and CA1 during state transitions. (A-L) DUPLEX recordings in V1. (A) *Left*, Representative raw epifluorescence (*top*) and activity-mask (*bottom*) images of a single field-of-view from a NDNF-*Cre*^*+*^*/*VIP-*Flp*^*+*^ mouse expressing Ace-mNeon2 in NDNF-interneurons (green) and pAce in VIP-cells (blue). Scale bar: 50 μm. (B) *ΔF/F* traces from the ROIs numbered in (A) aligned to air puff onset (vertical dashed line). Ace-mNeon2 traces are inverted for visualization purposes. Also shown (*top*), air puff onset, locomotory speed and pupil diameter. (C) Distribution of pairwise zero time-lag inter-type correlation coefficients, computed within a 2 s window before and 1 s window after air puff for all mice (n=908 NDNF-VIP pairs from 393 NDNF- and 188 VIP-cells, 7 mice). (D) Pairwise spike modulation indices for neighboring NDNF- and VIP-neurons for all mice. Each red dot represents values obtained for a single inter-type pair. (E-H) Same as (A-D) for DUPLEX recordings in SST-*Cre*^*+*^*/*VIP-*Flp*^*+*^ mice (n=242 SST-VIP pairs from 103 SST- and 79 VIP-cells, 6 mice). (I-L) Same as (A-D) for DUPLEX recordings in AAV-CaMKII-*Cre*/VIP-*Flp*^*+*^ mice (n=183 PN-VIP pairs from 75 PNs and 67 VIP-cells, 4 mice). (M-U) Dual color voltage recordings from hippocampal projection neurons. (M) Representative field-of-view of the red and green channels in gray scale and their overlaid spatial footprint showing 30 and 13 identified neurons, respectively. Scale bar: 50 μm (N) *ΔF/F* traces for all neurons in (M), aligned to the rest-run transition. Also shown (*top*), air puff onset and wheel speed. (O) Average mouse speed for 5 trials (black). Individual trials are shown in grey. (P) Average firing rate (250 ms window) for each projection-specific class. Shaded areas represent 95% CI. (Q) Firing rate fold change during a 2 s window before and after air puff. Shuffle is computed across random spike train permutations for each neuron. (R) Pairwise average coherence matrices during rest and run, computed for 1-50 Hz subthreshold range for across all neurons (n=135 neurons, 1 mouse). Note the increased average coherence during run versus rest, directly above the diagonal. (S-U) Average coherence within the (S) ACC-projecting subclass; (T) the EC-projecting subclass; and (U) across the two subclasses. Shaded areas represent 95% CI.

In contrast, in DUPLEX recordings from SST- and VIP-interneurons, the two cell types exhibited a more pronounced anti-correlation during state transitions compared to baseline conditions, when their activity was already negatively correlated (mean ± S.E.M. correlation coefficients were: −0.23 ± 0.02 before versus −0.45 ± 0.02 after air puff, *P*<0.0001, paired t-test) (Fig. 3, H and I, and 5, E-H). Our results suggest enhanced mutual inhibition during arousal when both populations exhibit stark increases in firing rates (Fig. 2, C and D). Further, pairwise distribution of spike modulation indices revealed stronger clustering in the top left quadrant, arising from a greater number of positively modulated SST-neurons amidst negatively modulated VIP-neighbors (Fig. 5H). Together, DUPLEX uncovers the millisecond-scale antagonism in the voltage dynamics between the two populations, even though major fractions of both neuron-types increase their spontaneous activity during state transitions (Fig. 2, C and D).

Lastly, while under baseline conditions, the activity of VIP-interneurons and excitatory cells were uncorrelated (*53*), arousal did not change their relationship dynamics (Fig. 5, I-L) (mean ± S.E.M. correlation coefficients were: −0.08 ± 0.02 before versus −0.005 ± 0.01 after air puff) (Fig. 5K). Further, pairwise distribution of spike modulation indices revealed no noticeable clustering, suggesting diverse spike rate modulation in VIP-cells and neighboring PNs for the targeted subclass (Fig. 5L).

### Dual color voltage imaging of hippocampal projection neurons

While DUPLEX can dissect activity patterns among non-overlapping populations, such as discrete excitatory and interneurons subtypes, where population overlap is unknown, subclasses can instead be demarcated by dual color voltage imaging using our spectrally orthogonal indicators. To this end, we used Ace-mNeon2 and VARNAM2 to investigate the extent to which adjacent glutamatergic neurons, targeting downstream neocortical brain regions, modulate their firing rates during a rest-to-run transition. To do so, we selectively expressed CaMKII-Ace-mNeon2 and CaMKII-VARNAM2 in anterior cingulate cortex (ACC)- and EC-projecting CA1 pyramidal neurons, respectively, using retrograde AAV labeling. We then performed dual color voltage imaging in an injected mouse, head-restrained on a running wheel, when brief air puff delivery elicited a rest-to-run behavioral state transition (Fig. 5M, see Methods).

We recorded from a total of 101 ACC-projecting and 34 EC-projecting neurons from 5 fields-of-view within the same animal, with a single field-of-view containing as many as 30 and 13 neurons of the two subpopulations, respectively (Fig. 5, M and N). Both Ace-mNeon2 and VARNAM2 captured the diversity of hippocampal intracellular dynamics, with isolated spikes, bursting activity and subthreshold oscillations, as previously reported (*20, 22*). We assessed whether ACC- and EC-projecting CA1 glutamatergic ensembles exhibited distinct supra- and subthreshold dynamics during state transitions (*6-9*).

Interestingly, ACC-projecting cells synchronously increased their mean firing rates during locomotion, whereas, EC-projecting neurons showed bimodal behavior, with some neurons exhibiting an increase and others showing a decrease in spike rate (log_2_ fold change (mean ± S.E.M.): 0.9 ± 0.1 versus 0.1 ± 0.2, *P*<0.001 and *P*=0.21 (versus shuffle), respectively) (Fig. 5, M-Q, and S21). We further observed an increase of subthreshold coherence during state transitions, suggesting an increase of excitatory/inhibitory synaptic inputs within neuronal subclasses. Interestingly, however, we found that each subclass synchronized at distinct theta frequencies (7.3 ± 0.3 Hz and 4.2 ± 0.2 Hz, respectively, *P*<0.001) (Fig. 5, R-U). These results support the existence of heterogenous parallel modules in the hippocampus (*56*).

### Simultaneous voltage imaging of three cell populations

Together, our sensors provide response polarity-based separation in addition to spectral unmixing of signals for dual population recordings. In principle, when combined, the four indicators are mutually compatible for concurrent voltage recordings from up to four distinct neuronal populations in intact networks. To demonstrate dual polarity and dual color multiplexing, we thus performed proof-of-concept voltage imaging from three distinct subtypes simultaneously.

In V1 of a SST-*Cre*^+^/VIP-*Flp*^+^ double transgenic mouse, we injected AAVs encoding floxed Ace-mNeon2 and *Flp*-dependent pAce, as well as a retrograde AAV expressing CaMKII-VARNAM2 in the AM to label a subpopulation of higher-order visual cortical projection neurons (Fig. 6A). We then performed simultaneous dual color voltage imaging under 1-photon microscopy to record from SST-, VIP- and excitatory cells concurrently, extracting their real-time voltage dynamics using Ace-mNeon2, pAce and VARNAM2, respectively, and inferring cell type based on both response polarity and indicator color (Fig. 6, B and C).

**Fig. 6.**
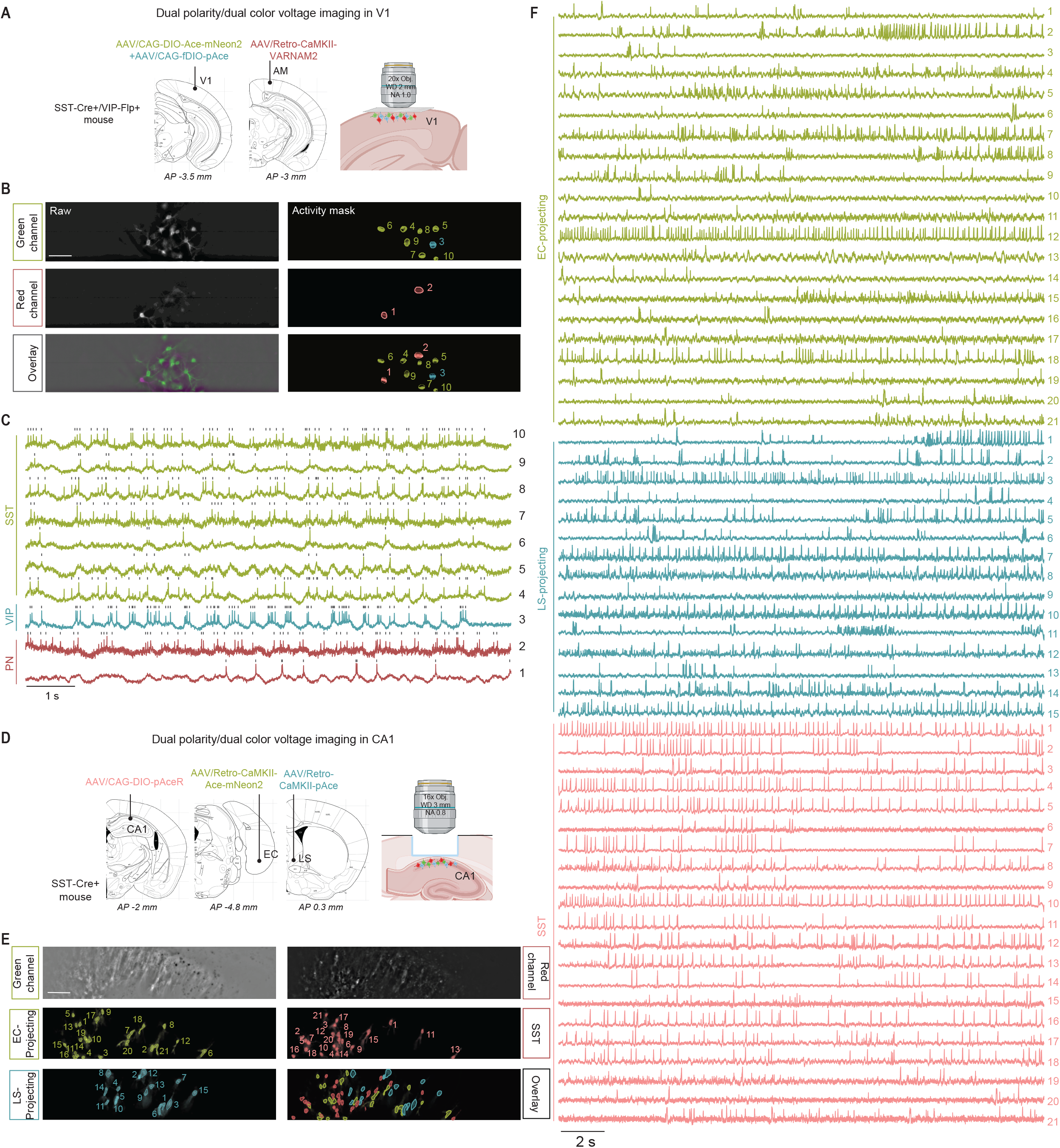
Simultaneous dual polarity and dual color imaging captures the voltage dynamics of three distinct cell classes in awake mice. (A) Schematic of AAV injections for three cell-type V1 imaging. (B) Raw epifluorescence (*left*) and activity-mask (*right*) images of a single field-of-view in the green and red channels, and overlay. Scale bar: 50 μm. (C) *ΔF/F* traces from the ROIs numbered in the mask overlay image in (B) and representing SST-interneurons, VIP-interneurons and PNs, expressing Ace-mNeon2 (green), pAce (blue) and VARNAM2 (dark red), respectively. Ace-mNeon2 and VARNAM2 traces are inverted for visualization purposes. (D) Schematic of AAV injections for three cell-type voltage imaging in CA1.(E) Representative field-of-view of the red and green channels with the respective spatial footprint of the identified neurons belonging to each of the three cell types. Scale bar: 50 μm. (F) *ΔF/F* traces for all neurons in (E), representing EC-projecting and LS-projecting excitatory neurons, and SST-interneurons expressing Ace-mNeon2 (green), pAce (blue) and pAceR (pale red), respectively. Ace-mNeon2 traces are inverted for visualization purposes.

Likewise, in the hippocampus, we retrogradely labeled EC-projecting and lateral septum (LS)-projecting CA1 pyramidal neurons, as well as local SST interneurons, using Ace-mNeon2, pAce and pAceR, respectively (Fig. 6D). Using dual color widefield imaging, we then captured the simultaneous spiking and subthreshold activity of as many as 57 neurons in a single field-of view belonging to the three neuronal subpopulations (21 EC-projecting, 15 LS-projecting and 21 SST-cells). All three ensembles belonged to distinct spatially or genetically non-overlapping populations (Fig. 6E), and all three indicators reported spiking at high signal-to-noise ratios (Fig. 6F).

## Conclusion

Our new suite of mutually compatible sensors enables high-speed voltage imaging of multiple genetically identified ensembles in awake animals, allowing real-time intra- and inter-population comparisons of voltage dynamics. Because our indicators are bright, they exhibit large signal-to-noise ratios even with inexpensive LED illumination, and being fully genetically encoded, they provide stable expression for long-term recordings without requiring exogenous dye applications (*57*).

Our indicators can provide large datasets for quantitative estimations of firing rates of targeted cell types in a biological paradigm. Our V1 data, acquired from ∼1200 cells using Ace-mNeon2, reveal that SST-interneurons exhibit a profound increase in spontaneous firing during arousal, recapitulating the results of targeted patch recordings (*4*). While our results do not support the VIP-neuron-centered PN disinhibition model (VIP ⊣ SST ⊣ PN), proposed based on Ca^2+^ imaging data (*2, 44, 58*), these differences may be context-dependent (*59*). That said, past interpretations of Ca^2+^ imaging data could be confounded by subtype-specific differences in intracellular Ca^2+^ dynamics. Further, intracellular Ca^2+^ release, evoked by state-dependent neuromodulatory action, might result in fluorescence responses which do not reflect ongoing activity (*59, 60*). Likewise, targeted whole-cell recordings underrepresent the contributions of sparse subpopulations (*e*.*g*. VIP-interneurons) (*2*). By comparison, GEVIs are poised to overcome such limitations of electrodes and Ca^2+^ imaging.

Traditionally, multipopulation imaging has been achieved using spectrally orthogonal indicators (*18, 48-51*), but, likely due to the limited set of mutually compatible sensors or the need for tailored optical hardware, simultaneous multichannel imaging is not widely performed in live animals. DUPLEX offers a simple alternative to distinguish two cell types using a single fluorescence channel. Here, DUPLEX unveiled a strong, time-varying antagonism between pairs of cortical SST and VIP interneurons that is further augmented by state transitions.

In the hippocampus, DUPLEX revealed subtype-specific subthreshold dynamics, which allows us to extract the relative contributions of select excitatory and inhibitory populations to the LFP frequency. While DUPLEX allows recordings from two genetically diverse subtypes, our spectrally compatible green and red indicators can be combined for dual population studies where mutual exclusivity is unknown. The ACC- and EC-projecting CA1 ensembles represent spatially distinct subpopulations, as seen from the non-overlapping expression of Ace-mNeon2 and VARNAM2 (Fig. 5M). Given the distinctive responses of these cell-types to state changes, they likely exhibit functional diversity as well.

Lastly, because our two polarity green indicators free up the spectral bandwidth, they can be further multiplexed with red-shifted sensors for simultaneous triple or quadruple population studies or can accommodate optogenetic control in the second channel. Collectively, our indicators empower current approaches to unravel intercellular interactions within and between targeted populations in intact networks.

## Acknowledgments

The authors thank the Pierce scientific staff C. Gardiner, M. Izydorsczak, P. O’Brien, X. Liu, T. Liu and R. O’Brien for technical assistance; the Pierce workshop members J. Buckley and A. Wilkins for instrumentation support; G. Lur, University of California, Irvine, and K. Ferguson, Yale School of Medicine, New Haven, for advice on V1 imaging; J. Verhagen, Pierce Laboratory and Yale School of Medicine, for comments on the manuscript. We further thank members of the Schnitzer laboratory J. Li for animal husbandry and genotyping, Y. Zhang for virus handling and surgical advice, G. Delamare for instrumentation support and F. Dinc and J. Li for computational consultations. All schematic representations in the manuscript were created with BioRender.com.

## Author contributions

MK and GV conceived the project, designed, screened and characterized the sensors, performed molecular biology and rAAV cloning; MK performed visual cortical surgeries; GV built the *in vivo* setup and wrote the software for V1 imaging; MK and GV conceived and performed *in vivo* V1 experiments and data analyses; SH conceived and performed the hippocampal experiments, wrote the *SpikeImagingAnalysis* pipeline and performed data analyses for CA1 studies; CH performed and analyzed the fly experiments; RC built the optical setup used for CA1 imaging; JL designed the transgenic flies; JAC shared the running wheel and provided advice on V1 imaging; MK and GV wrote the manuscript with contributions from all authors; MJS and VAP oversaw various aspects of the project.

## Data and materials availability

All sequence information will be deposited on NCBI, plasmids and rAAVs will be made available via Addgene and UNC Vector Core, fly stocks will be deposited at the Bloomington FlyBase and MATLAB codes will be shared on Github.

