## Supplementary information for "Dual polarity voltage imaging reveals subthreshold dynamics and concurrent spiking patterns of multiple neuron-types"

**This PDF file includes:**

Materials and Methods

Figs. S1 to S21

Tables S1

### Materials and Methods

**Plasmids.** For high-throughput screening in HEK cells, VARNAM and Ace-mNeon were inserted in a modified pCAGGS backbone (*18*). A nuclear localization sequence (NLS)-tagged FP reporter (mCerulean for VARNAM and mCherry for Ace-mNeon) was fused to the C-terminus of the indicator interspersed by a T2A ribosomal skipping signal. For *in vivo* voltage imaging, the indicators were cloned in adeno-associated viral (AAV) backbones. pAAV-CaMKII-Ace-mNeon2, pAAV-CaMKII-VARNAM2 and pAAV-CaMKII-pAce were cloned by inserting the respective indicator sequences in place of EGFP between the KpnI and HindIII sites of pAAV-CaMKII-EGFP (Addgene #50469). *Cre*-dependent constructs for Ace-mNeon2 and pAceR were generated by cloning the respective inverted open reading frame (ORF) sequences in pAAV-CAG-Flex-EGFP (Addgene #59331) by replacing the floxed EGFP cassette between the AscI and NheI sites. Likewise, *Flp*-dependent VARNAM2 and pAce constructs were cloned by introducing the respective inverted ORF sequences in place of the floxed mNeonGreen between the AscI and NheI sites in pAAV-CAG-fDIO-mNeonGreen (Addgene #99133). All AAV constructs above included a Golgi/ER export sequence and a Kv2.1 proximal restriction and clustering sequence (Lim et al, 2000, Neuron) at the C-terminus of the indicator sequence for improved membrane trafficking and perisomatic localization, respectively.

**Site-directed saturation mutagenesis.** Mutagenic libraries were generated on Ace-mNeon-T2A-NLS-mCherry and VARNAM-T2A-NLS-mCerulean backbones using a pre-established protocol (*18*). Briefly, a set of 4 forward primers, containing degenerate codons WKC, NMC, VWG or DGG at the target site, was pooled with a single, partially overlapping reverse primer and mutagenizing PCR reactions were set up using CloneAmp™ polymerase (Clontech). Following DpnI treatment to digest unmutated template, the linear PCR products were circularized using InFusion® ligase (Clontech) and transformed in TOP10 competent cells (Invitrogen). To obtain up to 19 unique AA substitutions at a single target site, 48 colonies were picked and cultured in 96 deep-well culture plates. Plasmid DNA was isolated using Nucleospin® 96 Plasmid kit (Macherey-Nagel) on an epMotion 5075 liquid handling workstation (Eppendorf), and purified DNA collected in 96-well plates. The libraries were sequenced after voltage screening to identify the individual mutations and ensure at least 19 variants were obtained at every site.

**Maintenance and transfection of HEK and excitable HEK cells.** HEK293 cells (ATCC® CRL-1573™) were maintained in high-glucose DMEM/10% FBS. Excitable HEK cells (ATCC® CRL-3269™) were maintained in DMEM/F12, 10% FBS, 1% penicillin (100 U $ml^{-1}$), streptomycin (100 µg $ml^{-1}$), geneticin (500 µg $ml^{-1}$), and puromycin (2 µg $ml^{-1}$) as described previously (*22*). ~24 h prior to transfection, the cells were plated on poly L-lysine-coated 12 mm coverslips or 96-well glass-bottomed plates in antibiotic-free media. Plasmid DNA (0.5 µg/12 mm coverslip or 0.2 µg/well of a 96-well plate) was transfected using Lipofectamine® (ThermoFisher), and cells were imaged 1-2 d post-transfection. Both cell lines served solely as a system for expression and characterization of voltage indicators rather than the subject of investigation.

**High-throughput semi-automated voltage screening.** High-throughput voltage screening was performed on a custom-built platform described previously (*18*). A white light source pE-4000 (CoolLED, UK) was used for sample illumination. For imaging Ace-mNeon, we used a 472/30 nm excitation filter, 495 nm dichroic mirror and 520/35 nm emission filter (Semrock). For VARNAM and mCherry (reporter fluorophore in Ace-mNeon constructs), we used a 560/40 nm excitation filter, 585 nm dichroic mirror and 630/75 nm emission filter (Chroma Technologies Corporation). The mCerulean reporter in the VARNAM constructs was imaged using a 455/40 nm excitation filter, 458 nm dichroic mirror and 480/30 nm emission filter (Semrock). After identifying transfected cells in the reporter channel, time-series fluorescence images were captured in the respective GEVI channels using ORCA Flash4.0 sCMOS camera (Hamamatsu) at 50 Hz. A single pulse of 60 V/0.5 ms was applied using a Grass S48 Stimulator, 1 s after baseline fluorescence acquisition ($F_0$). The percentage change in fluorescence at time *t* was obtained using the formula

$$\frac{\Delta F}{F_0}\% = \frac{F(t) - F_0}{F_0} x\ 100$$

Platform automation, sample illumination, data acquisition and parallel data analysis were controlled using custom-written virtual instruments in LabView®.

**Fluorescence voltage imaging in cultured HEK cells.** Whole-cell voltage clamp recordings were performed between 22-25˚C. The bath solution contained (in mM): 125 NaCl, 2 KCl, 10 HEPES, 30 glucose, 3 $CaCl_2$, 1 $MgCl_2$ (~310 mOsm $l^{-1}$, pH 7.3). The intracellular solution consisted of (in mM): 125 K-gluconate, 8 NaCl, 0.6 $MgCl_2$, 0.1 $CaCl_2$, 1 EGTA, 4 $Mg_2$-ATP, 0.4 Na-GTP and 10

HEPES. Patch pipettes had tip resistances of 4-5 MΩ, yielding series resistances <20 MΩ. In the voltage clamp mode, voltage steps in increments of 20 mV/ 0.5 s were applied via a MultiClamp amplifier controlled by pClamp10 software.

Fluorescence recordings were obtained using an Olympus upright microscope. Cells were visualized under a 1.0 NA 60x water-immersion objective lens. Ace-mNeon and pAce were imaged using a 505 nm light emitting diode (LED) (Thorlabs) and a filter set comprising 509/22 nm excitation filter, 526 nm dichroic mirror and 544/24 nm emission filter (Semrock). VARNAM and pAceR were illuminated using a 565 nm LED (Thorlabs), 560/40 nm excitation filter, 585 nm dichroic mirror and 630/75 nm emission filter (Semrock). The illumination intensity at the sample plane for all indicators was 15-20 mW $mm^{-2}$. Fluorescence time-series images (1-5 kHz) were captured using a NeuroCCD camera, controlled via NeuroPlex software (RedShirtImaging, GA).

**Intracranial V1 injections and cranial window implants.** Animal experiments were performed according to the guidelines of the National Institutes of Health and approved by The John B. Pierce Laboratory Animal Care and Use Committee.

C57BL/6J, VIP-Cre, SST-Cre, VIP-FlpO and NDNF-Cre lines were purchased from The Jackson Laboratory (JAX #664, #31628, #13044, #28578, #28536). NDNF-$Cre^{+/+}$ or SST-$Cre^{+/+}$ mice were crossed with VIP-$FlpO^{+/+}$ to generate the NDNF-$Cre^{+}$/VIP-$FlpO^{+}$ and SST-$Cre^{+}$/VIP-$FlpO^{+}$ double driver lines. The following AAV vectors were custom-produced at the University of North Carolina Vector Core at high titers (>1 x $10^{13}$ GC $ml^{-1}$): AAV/DJ-CAG-DIO-Ace-mNeon2, AAV2/Retro-CaMKII-Ace-mNeon2, AAV/DJ-CAG-fDIO-pAce, AAV2/Retro-CaMKII-pAce, AAV/DJ-CAG-fDIO-VARNAM2, AAV2/Retro-CaMKII-VARNAM2 and AAV/DJ-CAG-DIO-pAceR. All AAV vectors above encode soma-targeted versions of the indicators. AAV5-CaMKII-mCherry-Cre and AAV2/Retro-EF1A-mCherry-IRES-Cre viruses were purchased from the UNC Vector Core and Addgene (#55632), respectively.

Mice were used without regard to sex. Stereotactic AAV injections were performed in 3-4 month-old mice under 1.5% (v/v) isoflurane anesthesia. The coordinates for V1 injections were (in mm from Bregma; AP, ML): 2.5, 2.5; 2.8, 2.0; 3.0, 2.5; 3.4, 2.0; 3.6, 2.75; DV=0.30. For projection labeling in the anteromedial (AM) visual cortical area, the coordinates were (AP, ML, DV): 3.0, 0.96, 0.40. Injections were performed using beveled glass micropipettes delivering a total of 50-100 nl virus/site at a rate of 100 nl/min. For DUPLEX imaging, AAV/DJ-CAG-DIO-

Ace-mNeon2 and AAV/DJ-CAG-fDIO-pAce viruses were mixed at 1:3 and 1:1 (v/v) ratios for injections in NDNF-Cre$^+$/VIP-FlpO$^+$ and SST-Cre$^+$/VIP-FlpO$^+$ mice, respectively. For dual color imaging, AAV/DJ-CAG-DIO-Ace-mNeon2 and AAV/DJ-CAG-fDIO-VARNAM2 were mixed at a 1:1 (v/v) ratio for injections in a SST-Cre$^+$/VIP-FlpO$^+$ background. For sparse labeling of V1 PNs, AAV/DJ-CAG-DIO-Ace-mNeon2 was mixed with AAV5-CaMKII-mCherry-Cre at a 2:1 (v/v) ratio. Alternatively, the AAV/DJ-CAG-DIO-Ace-mNeon2 virus alone was injected locally, while the AAV2/Retro-EF1A-mCherry-IRES-Cre virus was delivered to the AM cortical area.

Headpost and cranial window were implanted between 25-30 days post-injection with some modifications to a pre-established protocol (*61*). Mice were anesthetized using a mixture of ketamine (100 mg/kg) and xylazine (16 mg/kg). A custom-designed titanium headpost was affixed to the skull using C&B Metabond (Parkell), following which a 3 mm diameter circular craniotomy was made over the injection area in V1. A sterile double-glass window, assembled from 1x 3 mm and 1x 5 mm diameter circular coverslips (Warner), was secured in place using Vetbond, as previously described. C&B Metabond was applied over the window margins and the remainder of the exposed skull. Carprofen (5 mg/kg, SC, every 24 h for 48 h) and buprenorphine (100 µg/kg, SC, every 12 h for 48 h) were administered as part of postoperative analgesia. Prior to awake imaging experiments, headposted mice were handled and habituated to run on a custom wheel every day for up to 2 weeks in 10-20 min training sessions.

**Mouse hippocampal surgeries.** The Stanford Administrative Panel on Laboratory Animal Care (APLAC) approved all procedures involving animals, and we complied with all ethical regulations. C57BL/6J and SST-Cre lines were purchased from The Jackson Laboratory (JAX #664 and #013044). Mice (aged 8–16 weeks at start) underwent two surgical procedures under isoflurane anesthesia (1.5%–2% in $O_2$). In the first procedure, we injected AAVs to express the fluorescent voltage indicators. In the second procedure, we inserted a cannular implant for hippocampal imaging. The coordinates for CA1 injections were (in mm from Bregma; AP, ML, DV) -1.8, -2.5, -1.1 and-1.3, -1.3, -1.1. The coordinates for retrograde labeling in the lateral septum (LS), entorhinal cortex (EC) and anterior cingulate cortex (ACC) were, respectively, (in mm from Bregma; AP, ML, DV): 0.3, 0.3, -3; -4.8, 3.3, -3.5 and 0.8, 0.2, -1.5. We used the following viruses: AAV2/DJ-Retro-CaMKII-VARNAM2 and AAV2/DJ-Retro-CaMKII-Ace-mNeon2 to label EC-projecting and ACC-projecting neurons, respectively (Fig.5); AAV2/DJ-CAG-DIO-Ace-mNeon2 and AAV2/DJ-Retro-CaMKII-pAce to label SST neurons and EC-projecting neurons, respectively

(Fig.4); and AAV2/DJ-Retro-CaMKII-Ace-mNeon2, AAV2/DJ-Retro-CaMKII-pAce and AAV2/DJ-CAG-DIO-pAceR to label EC-projecting and LS-projecting neurons, SST-neurons, respectively (Fig.6). All viral vectors had titers of $>10^{12}$ GC $ml^{-1}$ and were custom-produced at the UNC Vector Core. We performed cannular implantation ~1 week post-injection. We scored the skull manually to remove the connective tissue. After cleaning the skull with $H_2O_2$ and rinsing with Ringer's, we drilled a 3 mm opening above the dorsal CA1 (right hemisphere), large enough for the cannular implant to fit in snugly. The cortex above the dorsal CA1 was aspirated with a *30G* blunt needle and rinsed with ice-cold Ringer's. Cortical aspiration was continued until the light-scattering myelinated axon bundles of the corpus callosum became visible. At this point, the top layers were gently removed, while making sure to leave the *stratum oriens* intact. Mice with damaged CA1 were discarded, as a damaged hippocampus is prone to local epileptic seizure. The implant was then inserted and affixed to the skull using UV-curable adhesive (Loctite 3105). Carprofen (5 mg $kg^{-1}$) was administered 30 min prior to the end of the surgery to mitigate pain. A custom stainless steel head bar was secured to the mouse's skull, enabling head-fixation during *in vivo* imaging. The overall implant was secured in place with blue light-cured resin (Flow-it ALC, Pentron). Carprofen (5 mg $kg^{-1}$) was administered up to two days post-surgery as part of postoperative analgesia. Mice recovered for ~2 weeks prior to imaging.

**Implant fabrication, surgery and LFP recordings.** Cannular implant fabrication was performed under a stereomicroscope (Leica MZ7-5) on a fully equipped solder station. The hippocampal cannula was made of a round borosilicate cover glass (3 mm diameter, 170 µm ± 5um thickness, Schott) glued to a 3 mm OD, 1.5 mm long stainless-steel ring, using UV-curable adhesive (Loctite 3105). For the dual modality optical-LFP hippocampal recordings, we prepared an optrode using three 0.002" tungsten wires (99.95% CS, with a single polyimide insulation, M215580, California Fine Wire) positioned at three different axial positions for redundancies. Tungsten wires were glued in place onto the optical cannula using UV light-cured adhesive. Next, tungsten wires were soldered into a gold-plated pin head (0.031", WPI). The reference channel was inserted into the cerebellum and made of coated stainless-steel wire (0.005" bare, 0.008" coated, 100 feet, A-M systems). Separately, we fabricated a connector using the gold-plated sockets (0.031", WPI) and a connector board (EIB8, Neuralynx). We pre-amplified the signals using a digital recording head-stage (RHD 16-Channel, Part #C3334, Intantech). We amplified electrophysiological signals using a USB acquisition board (RHD USB interface board, Part #C3100, Intantech). Electrical LFP signals were sampled at 2 kHz. To synchronize the optical and electrical recording modalities, we used a random TTL (syncTTL) externally and independently generated using a National Instruments Daq system (LabView interface). To homogenize the two sampling rates (600 Hz and 2 kHz, respectively), we first band-pass filtered the LFP traces (filter's high-frequency 3 dB cutoff frequency was 500 Hz and the low-frequency 3 dB cutoff ranged between 0.1 Hz). Then, we interpolated the LFP traces and the syncTTL to match the optical frame rate. Finally, the optical and electrical traces were aligned using the independently recorded syncTTL signal.

**Behavioral paradigm to study rest-to-run transitions in the CA1.** The custom mouse wheel consisted of a 3D-printed PLA plastic (8 cm wide, 13 cm diameter). The wheel surface was covered with self-adherent wrap (Coban, 3M). Angular displacement was tracked using a rotary encoder (Optical AB Phase Quadrature Encoder 600P/R, Amazon). A data acquisition device with LabView GUI (BNC-2090A, National Instruments) was used to register wheel displacement. Mice were habituated on the wheel for at least 2 days, for at least 15 min per day, before imaging sessions began. To trigger rest-to-run behavioral state transitions, brief air puff was delivered, directed to the mouse's back. The air puff intensity was ~30 psi and was administered via a 20-gauge blunt needle placed 1-2 cm away from the mouse. High-speed voltage imaging of a 30 s trial per field-of-view (18,000 frames at 600 Hz) was performed starting at rest. A TTL (5 pulses at 5 Hz) triggered the air puff after a >10 second baseline. Each experimental session consisted of 6 fields-of-view with 1 trial per field-of-view.

***In vivo* voltage imaging in the primary visual cortex.** Prior to awake imaging experiments, headposted mice were handled and acclimated to a custom 15 cm diameter 3 D-printed wheel for ~2 weeks in 10 min training sessions/day.

For *in vivo* neocortical voltage imaging, mice implanted with cranial windows were head-fixed by securing the headpost to two points on the custom wheel via thumb screws. Neurons expressing soma-targeted indicators were imaged on a custom-built upright fluorescence microscope using a 20X 1.0 NA Olympus XLUMPLFLN water immersion objective. Excitation light from a 505 nm or 565 nm LED (M505L4 or M565L3, Thorlabs) was collimated using an aspheric condenser lens (ACL2520U-A) and attached to an epi-illuminator module (WFA2001, Thorlabs). An EYFP filter set (49003-ET-EYFP, Chroma), comprising a 500/20 excitation filter, 515 nm LP dichroic mirror and 535/30 nm emission filter, was introduced into the optical path for

Ace-mNeon and pAce. An mCherry filter set (49008-ET-mCherry, Chroma), consisting of a 560/40 excitation filter, 585 nm LP dichroic mirror and 630/75 nm emission filter, was used for fluorescence imaging with VARNAM and pAceR. Light intensity at the sample plane was ~25 mW mm$^{-2}$. A 0.75X camera tube (WFA4101, Thorlabs) was used to focus the image onto a high speed sCMOS camera (Orca Flash 4.0 v3, Hamamatsu). Images were collected at 400 Hz with 4x binning. A second infrared camera (Basler A602f) was used to measure pupil diameter at a frame rate of 15 Hz.

Visual stimuli were generated using the Psychophysics toolbox on MATLAB and were presented to awake, stationary mice on an LCD monitor (1280 x 1024 pixels, 20 x 16 inches, 60 Hz refresh rate, mean luminance of 250 cd m$^{-2}$). The monitor was placed at ~15 cm from the animal, perpendicular to the surface of the right eye (contralateral to injection area). To pre-map the receptive field center of an imaged neuron, gratings were presented in a 3 x 4 grid, covering the entire screen. To obtain orientation-selective responses, drifting sinusoidal gratings were presented at eight orientations, separated by 45°. The spatial frequency was set at 0.04 cycles per division and temporal frequency at 2 Hz. Trials lasted 16 s with stimulus periods of 500 ms and interstimulus intervals of 500 ms.

To induce a state change, a brief 50 ms air puff was delivered to mice at rest using a small tube aimed at the back of the animal. Wheel position was tracked using a programmable angle sensor (Digikey). A data acquisition device (NI USB-6259, National Instruments) was used to register wheel position and deliver TTL pulses to initiate image acquisition. A custom-written code (LabView) controlled data acquisition.

Pupil diameter was determined in real-time using the edge-detection functionality in the LabView Vision Development module. An annulus region-of-interest was drawn over the pupil, with an inner circle located at the center of the eye and an outer circle extending past the edge of the pupil. Dark to light transition edges were detected as lines emerging from the inner to the outer circle. A circle was fit based on the identified edges to calculate the pupil diameter.

**Optical setup for *in vivo* dual color recordings in CA1.** For *in vivo* imaging in CA1, we used a 16X Nikon CFI LWD Plan Fluorite objective lens (0.80 NA, 3.0 mm WD, water immersion). Excitation light from 470 nm and 545 nm LEDs was combined using a LP dichroic. In the emission path, we used a 550 nm SP dichroic mirror and an emission filter specific for each color channel:

520/41nm for Ace-mNeon2 and pAce and 609/62 nm for VARNAM2 and pAceR. Light intensity at the sample plane was ~25-50 $mW/mm^{-2}$. High-speed videos were captured using Hamamatsu ORCA Fusion Digital CMOS camera (Hamamatsu Photonics K.K., C14440-20UP). A custom-written LabView GUI controlled data acquisition. Images were collected at 600 Hz with no binning. Live data were stored as DCAM image files (.dcimg), and batch-processed offline using custom-written *SpikeImagingAnalysis* pipeline (MATLAB, Mathworks).

**V1 data acquisition and analysis.** Data analysis was performed using a python-based voltage imaging data analysis package (VolPy) in Caiman (*62*). Briefly, the video file from each trial was motion-corrected using the NoRMCorre algorithm followed by automated region-of-interest (ROI) identification using the Mask R-CNN neural network architecture. Mask R-CNN uses a pre-trained convolutional network (ResNet) and a Feature Pyramid Network (FPN) backbone to extract spatial features. A Regional Proposal Network identified initial bounding boxes. The spatial features and initial bounding boxes were further refined and used to create binary masks for trace extraction.

We made several modifications to the VolPy code to be better compatible with our data files and for analyses of the dual polarity recordings. First, the code was modified to support the native .dcimg file format of data acquired using the Hamamatsu camera. Next, while VolPy requires user input to identify the polarity of the traces in each recording, we included additional lines of code to identify the polarity in DUPLEX recordings. This was done by calculating the number of spikes in both directions in the optical trace that exceeded a threshold of 3 x S.D. and assigning the polarity to that direction which contained the largest number of spikes that crossed the threshold.

Spike timing and subthreshold activity were computed using the SpikePursuit algorithm as part of the same VolPy package. The data from VolPy were exported as MATLAB files for further downstream analyses. The approximate firing rate (FR) was determined by sliding a rectangular window along the spike train with $\Delta t$=100 ms and smoothed with a Gaussian kernel using the *smooth* function in MATLAB. The air puff-induced spike modulation index was calculated as $(FR_{arousal} - FR_{baseline})/(FR_{arousal} + FR_{baseline})$, where FR is the firing rate during a 750 ms time window under baseline conditions and upon arousal, and their statistical significance was evaluated Wilcoxon rank sum test. Peristimulus time histograms for cells with significant increase or decrease in air puff-induced firing rates were plotted by aligning the spike rates to stimulus onset. The correlation coefficients and cross-correlation values were calculated from the *ΔF/F* traces using the corrcoef and xcorr functions in MATLAB.

For subthreshold power calculations, the subthreshold traces obtained using the SpikePursuit algorithm were normalized by their respective spike amplitudes from the *ΔF/F* traces and were used to estimate the power spectral density during a 3 s window before and after air puff delivery.

**Hippocampal data analyses - *SpikeImagingAnalysis* Pipeline.** Hippocampal data were analyzed using a custom analysis pipeline written in MATLAB (*SpikeImagingAnalysis*, https://github.com/sihaziza/SpikeImagingAnalysis_public). Our pipeline consists of 5 steps: (i) loading, (ii) motion correction, (iii) detrending, (iv) demixing and (v) spike timing estimation. (i) Raw .dcimg data were loaded, binned 2x2 and saved as .h5. (ii) Movies were motion-corrected using NoRMCorre and cropped to remove NaN borders. The NoRMcorre parameter shift_method was set to ‘cubic’, instead of the default ‘fft’. (iii) Detrending consisted of pixel-wise normalization by the time-varying baseline fluorescence $F_0$, estimated using a pixel-wise temporal low-pass filter ($4^{th}$ order Butterworth filter, 0.5 Hz cut-off frequency). This method was found to be faster and more accurate than previously reported approaches (spline or exponential fit). (iv) Demixing of neural units to find the spatial footprint and temporal trace of neurons was performed using EXTRACT (*63*). For DUPLEX experiments involving spectrally overlapping Ace-mNeon2 and pAce, EXTRACT was computed two times: once on the detrended movie (+$\Delta F/F_0$) and once on the flipped detrended movie (-$\Delta F/F_0$). (v) To precisely estimate spike bursts, typically observed for hippocampal neurons, we used the statistical method MLspikes (*64*). It requires five input parameters: saturation **S**, single-spike amplitude **A**, single-spike time constant **tau**, baseline fluctuation **drift** and noise level **sigma**. We set the saturation parameter to **S**=1, defined as the number of spikes to reach full amplitude. **A** and **tau** were estimated by a double exponential fit on the spike-triggered average, computed on isolated spikes. **Drift** was estimated as peak-to-peak signal on the low-pass-filtered time trace ($4^{th}$ order Butterworth filter, 50 Hz cut-off frequency). **Sigma** was estimated as the standard deviation of the residual between the raw and denoised trace (wavelet denoising of degree one).

**Suprathreshold data analyses.** For a given field-of-view, all traces (spike rasters and speed trace) were centered around the rest-to-run transition, detected as a speed change (thresholded at >2 cm/s

± 7.5 s). Data from all neurons across all fields-of-view were consolidated to compute the time-varying firing rate (sliding window of 100 ms) and the rest-to-run fold change of instantaneous firing rates (2 s pre- and post-state transition). For a given cell class, the time-varying firing rate was averaged across all neurons.

**Subthreshold data analyses.** All neurons across all fields-of-view were centered around the rest-to-run transition. Pairwise coherence matrices were constructed for both states using the corresponding low-pass filtered versions of the baseline output of MLspikes ($4^{th}$ order Butterworth filter, cut-off frequency of 50 Hz). To quantify the coherence noise level, pairwise coherence of neurons belonging to two independent fields-of-view was considered. For neuron-LFP coherence analysis, pair-wise coherence was computed between the baseline activity of the neurons and the LFP.

**Spike-Subthreshold-LFP cross-analysis.** To calculate the phase of spiking at the theta frequency (5–10 Hz), we first band-pass-filtered both the optical voltage trace and the LFP trace at theta frequency ($4^{th}$ order Butterworth filter). We then computed the phase for each filtered trace using a Hilbert transform. We calculated the timing of each spike relative to the period of that oscillation cycle in degrees.

**Statistics.** Statistical tests were performed using standard MATLAB functions. For two-sample comparisons of a single variable, a two-sided Student's t-test (paired or unpaired) was used, if normality test was valid. Otherwise, a non-parametric test was used (Mann-Whitney Wilcoxon test). The experiments were not randomized, and the investigators were not blinded to the experimental condition. Power analysis was not conducted prior to the experiments. No statistical methods were used to estimate sample sizes for the mouse studies. These numbers reflect our prior experience or what is reported in the literature. Data exclusion criteria were not pre-established, however, recordings with no spiking neurons were excluded.

**Fly stocks.** To create transgenic flies carrying the four improved FRET-opsin GEVIs, we synthesized codon-optimized transcripts for Ace-mNeon2, VARNAM2, pAce and pAceR using a commercial gene synthesis service (GenScript), and then inserted them in place of GFP in the *pJFRC7-20×UAS-IVS-mCD8::GFP* (Addgene #26220) and *pJFRC19-13×LexAop2-IVS-myr::GFP* backbones (Addgene #26224). After verifying the sequences of the plasmids, we inserted the plasmids into the attP40 or VK00027 phiC31 docking site to generate *20×UAS-Ace-*

*mNeon2*, *13×LexAop-Ace-mNeon2*, *20×UAS-pAce*, *13×LexAop-pAce*, *20×UAS-VARNAM2*, *13×LexAop-VARNAM2*, *20×UAS-pAceR* and *13×LexAop-pAceR* flies using a commercial transformation service (Bestgene Inc.).

We obtained MB058B, MB296B, and MB085C split-GAL4 lines from the FlyLight team at Janelia Research Campus, and *R82C10-LexA* (#54981) from the Bloomington Stock Center. We raised all flies on standard cornmeal agar media under a 12 h light/dark cycle, at 25°C and 50% relative humidity.

**Dissection of live flies**. We created an optical window on the fly head using an ultraviolet (UV) laser microsurgery system, as described previously (*57*). In brief, we anesthetized the flies by placing them on ice for 1 min and then transferring them to a cooled surface (∼4 °C) consisting of an aluminum thermoelectric cooling block. We then glued the fly's thorax with a 125 μm-diameter fused silica optical fiber (PLMA-YDF-10/125, Nufern) on a custom-made plastic fixture. To minimize head motion, we glued the fly's head to the thorax using a UV light curing epoxy (NOA 89, Norland). After transferring the mounted fly to the surgery station, we created an optical window in the cuticle by laser-drilling a 150-μm-diameter hole (30–4¬0 pulses, delivered at 100 Hz, 36 mJ per pulse, as measured at the specimen plane). Immediately after surgery, we applied 1 mL of UV epoxy (NOA 68, Norland; Refractive index: 1.54; Transmission 420–1000 nm: ~100%) and cured it for ~30 s to seal the cuticle opening.

***In vivo* voltage imaging in flies**. To image voltage dynamics, we used a custom-built upright epi-fluorescence microscope and a 1.0 NA 20X water-immersion objective (XLUMPlanFL, Olympus). For Ace-mNeon and pAce imaging, we used a 503/20 nm excitation filter (Chroma), 518 nm dichroic (Chroma) and 534/30 nm emission filter (BrightLine). We illuminated the sample using the 500-nm wavelength module of the solid-state light source (Spectra X, Lumencor) with 5–7 mW of optical power at the specimen plane. For VARNAM and pAceR imaging, we used a 559/25 nm excitation filter (Semrock), 580 nm dichroic (Chroma) and 630/50 nm emission filter (Semrock). We illuminated the sample using the 550-nm wavelength module of the solid-state light source (Spectra X, Lumencor) with 8–10 mW of optical power at the specimen plane. We acquired images at 1000 Hz, using a scientific-grade CCD camera (Zyla 4.2, Andor) and 2 × 2 pixel binning.

To image odor-evoked activity, we perfused odors to the flies' antennae using a custom-built olfactometer that delivered constant airflow (200 mL/min) to either the control path (mineral oil) or the odor path (odorant dissolved in mineral oil). We presented 5% Isoamyl acetate (IAA; CAS# 123-92-2, Sigma-Aldrich Inc.), 3% Benzaldehyde (BEN; CAS# 100-52-7, Sigma-Aldrich Inc.) or apple cider vinegar (ACV; Bragg Inc.) to the fly through a probe needle (1.7 mm ID, Grainger Inc.), ~3 mm in front of the antenna. Every trial of odor delivery lasted for 5 s.

To analyze the voltage traces, we motion-corrected the raw videos using the Turboreg algorithm for image registration (http://bigwww.epfl.ch/thevenaz/turboreg/). We then selected pixels whose mean fluorescence intensity was in the top 10%; we defined their union as the region-of-interest (ROI), over which we computed spatially averaged, time-dependent changes in relative fluorescence intensity, $\Delta F_{(t)}/F_0$, where $F_0$ was the mean fluorescence in the ROI averaged over the entire video. We corrected for photobleaching by fitting a double exponential function to the mean fluorescence trace, $F_0$, and then normalizing $F_0$ by the fitted double exponential trace. To identify individual spikes, we high-pass filtered the $\Delta F_{(t)}/F_0$ trace by subtracting a median-filtered (40 ms window) version of the trace and then identified the spikes as local peaks that surpassed the threshold (>3 S.D. for PPL1-DANs, >2 S.D. for MBON-$\gamma$1pedc>α/β). We determined the approximate firing rate by sliding a rectangular window along the spike train with $\Delta t$ = 100 ms. To analyze spike waveforms, we determined the mean waveform by averaging across all spikes within a trial. We then performed a spline interpolation (10 μs intervals) of the mean waveform and, from the output, determined the spike amplitude and the spike duration. To compute spike detection fidelity d′, we used a signal detection framework that accounts for both the duration and intensity of fluorescence waveforms in optical recordings of action potentials (*17, 34*).

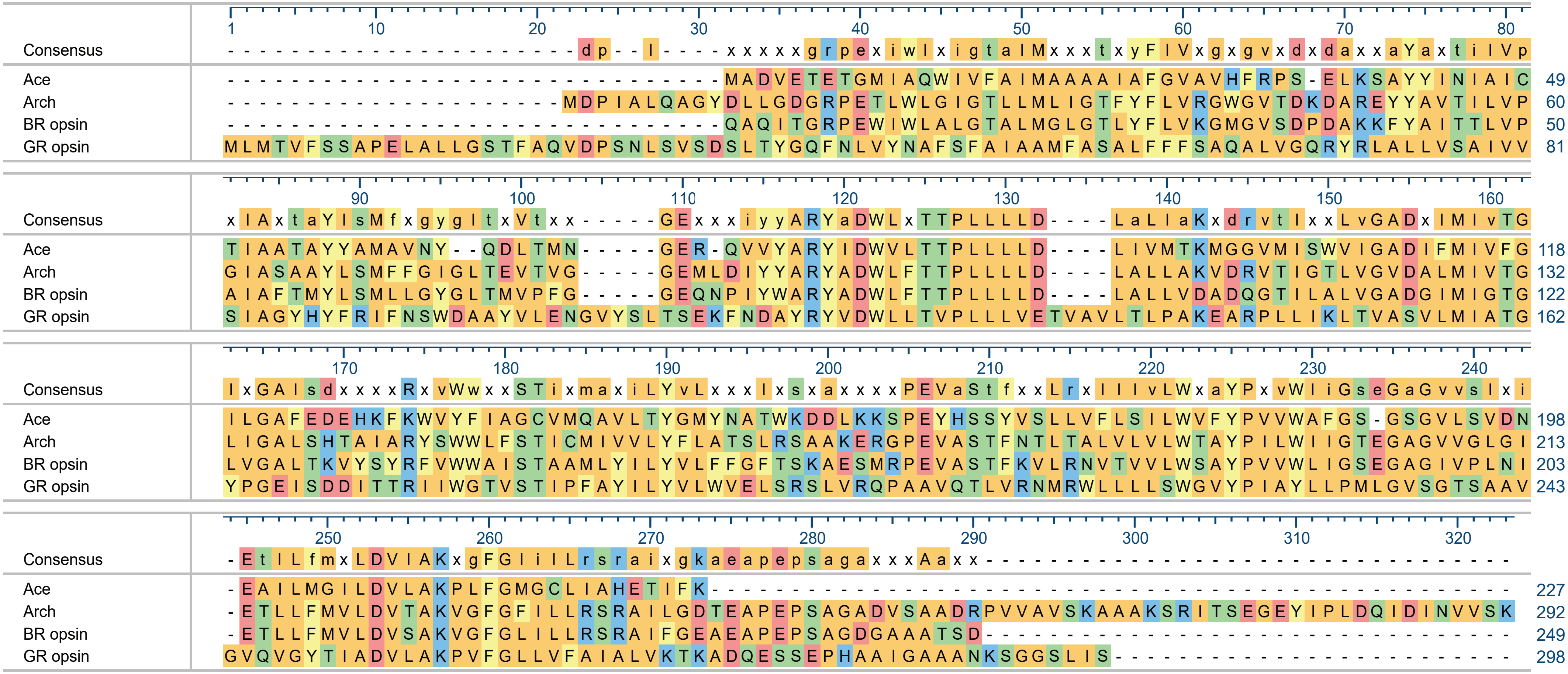


**Figure S1** | Alignment of amino acid sequences of *Acetabularia* rhodopsin II (Ace), Archaerhodopsin (Arch), bacteriorhodopsin (BR) and *Gloeobacter* rhodopsin (GR). Boxes indicate residues targeted for site-directed saturation mutagenesis. AAs are color-coded based on side-chain chemistry. Conserved residues are indicated in upper case.

Ace N81X mutagenesis in Ace-SY-mNeon

| | 1 | 2 | 3 | 4 | 5 | 6 | 7 | 8 | 9 | 10 | 11 | 12 |
|---|---|---|---|---|---|---|---|---|---|---|---|---|
| A | -10<br>N81H | -3<br>N81I | 0<br>N81A | -4<br>N81L | -2<br>N81P | -4<br>N81P | -3<br>N81N | -4<br>N81P | 0<br>N81I | -5<br>N81N | -6<br>N81R | -8<br>N81S |
| B | -3 | -2 | -3 | -8 | -2 | -4 | -8 | -4 | -2 | -12<br>N81A | -10 | -3 |
| C | -7 | 0 | -18<br>N81S | 0 | -11 | -3 | -10 | -10 | -9 | -5 | -14<br>N81S | 0 |
| D | -14<br>N81F | -2 | -16<br>N81S | -8 | -8 | -7 | -11 | -3 | -5 | -11<br>N81S | -6 | Control |

Ace-FP linker mutagenesis in Ace-mNeon 81S (well C3 from above) (G229X)

| | 1 | 2 | 3 | 4 | 5 | 6 | 7 | 8 | 9 | 10 | 11 | 12 |
|---|---|---|---|---|---|---|---|---|---|---|---|---|
| A | -25<br>G229G | -16<br>G229P | -13<br>G229E | -14<br>G229P | -15 | -18<br>G229E | -17<br>G229S | -18 | -20<br>G229H | -25<br>G229K | -17 | -16 |
| B | -17 | -15 | -17 | -18<br>G229T | -12 | -15 | -12 | -10 | -21<br>G229F | -12 | -13 | -18 |
| C | -14 | -13 | -10 | -24<br>G229F | -10 | -15 | -11 | -5 | -18<br>G229F | -14 | -16 | -8 |
| D | -16 | -17 | -6 | -12 | -12 | -8 | -9 | -9 | -18<br>G229F | -34<br>G229Y | -5 | -25<br>G229W |

Ace-FP linker mutagenesis in VARNAM Δ229 Δ230 (G231X)

| | 1 | 2 | 3 | 4 | 5 | 6 | 7 | 8 | 9 | 10 | 11 | 12 |
|---|---|---|---|---|---|---|---|---|---|---|---|---|
| A | -17<br>G231S | -26<br>G231I | -14<br>G231D | -22<br>G231T | 0<br>G231Q | -16<br>G231A | -10<br>G231E | -18<br>G231M | -8<br>G231P | -19<br>G231K | -16<br>G231A | -16<br>G231S |
| B | -20<br>G231L | -16<br>G231D | -18<br>G231M | -15<br>G231Q | -18<br>G231C | -15<br>G231T | -11<br>G231A | -18<br>G231F | -20<br>G231L | -17<br>G231R | -12<br>G231G | -23<br>G231T |
| C | -16<br>G231S | -21<br>G231K | -8<br>G231P | -12<br>G231D | -15<br>G231A | -17<br>G231C | -15<br>G231F | -20<br>G231K | -18<br>G231L | -15<br>G231E | -23<br>G231L | -18<br>G231N |
| D | -20<br>G231Q | -20<br>G231K | -17<br>G231M | -18<br>G231L | -17<br>G231S | -16<br>G231W | -17<br>G231F | -19<br>G231F | -18<br>G231M | -15<br>G231G | -16<br>G231E | Control |

**Figure S2** | High-throughput screening for voltage sensitivity of Ace-mNeon2 and VARNAM.
(A) 48 variants containing saturation mutations at Ace N81 on the backbone of Ace-mNeon were each transfected in electrically excitable HEK cells[25,26] and screened for voltage sensitivity on the high-throughput platform[18]. The maximum %Δ*F/F* obtained for each variant across 4 independent rounds of screening are indicated in each well together with the sequence information, where available. Color density corresponds to the size of the signal in the negative direction.
(B) Same as (A) for Ace-FP linker mutagenesis on Ace-mNeon 81S backbone.
(C) Same as (A) for Ace-FP linker mutagenesis on VARNAM backbone.

Ace opsin

ATGGCTGACGTGGAAACCGAGACCGGCATGATTGCACAGTGGATTGTCTTTGCTATTATGGCTGCTGCTGCTATTGCTTTTGGAGTGGCTGTGCACTTTC
M A D V E T T G M I A Q W I V F A I M A A A A I A F G V A V H F

Ace opsin

GGCCTTCAGAGCTGAAGAGCGCATACTATATCAACATTGCCATCTGCACTATCGCCGCTACCGCTTACTATGCAATGGCCGTGAACTACCAGGACCTGAC
R P S E L K S A Y Y I N I A I C T I A A T A Y Y A M A V N Y Q D L T

Ace opsin

AATGAATGGTGAAAGGCAGGTGGTCTACGCAAGATATATTAGCTGGGTGCTGACCACACCACTGCTCCTGCTCGATCTCATCGTCATGACCAAGATGGGC
M N G E R Q V V Y A R Y I S W V L T T P L L L L D L I V M T K M G

Ace opsin

GGAGTGATGATTTCTTGGGTCATCGGCGCAGACATTTTCATGATCGTGTTTGGTATTCTGGGCGCCTTCGAGGATGAACACAAGTTCAAATGGGTGTACT
G V M I S W V I G A D I F M I V F G I L G A F E D E H K F K W V Y

Ace opsin

TTATCGCTGGATGTGTGATGCAGGCAGTCCTGACATACGGGATGTATAACGCCACTTGGAAAGACGATCTGAAGAAAAGCCCCGAGTACCATAGCTCCTA
F I A G C V M Q A V L T Y G M Y N A T W K D D L K K S P E Y H S S Y

Ace opsin

TGTCAGTCTGCTCGTCTTCCTGTCAATCCTCTGGGTGTTTTATCCTGTCGTGTGGGCTTTCGGGTCTGGTAGTGGCGTGCTGTCCGTCGACAATGAGGCC
V S L L V F L S I L W V F Y P V V W A F G S G S G V L S V D N E A

Ace opsin | Linker | mNeonGreen

ATTCTCATGGGAATCCTGGATGTGCTCGCTAAGCCACTGTTTGGAATGGGGTGCCTCATTGCCCATGAGACTATCTTCAAGTCCTACCCAGCGACACATG
I L M G I L D V L A K P L F G M G C L I A H E T I F K S Y P A T H

mNeonGreen

AGTTACACATCTTTGGCTCCATCAACGGTGTGGACTTTGACATGGTGGGTCAGGGCACCGGCAATCCAAATGATGGTTATGAGGAGTTAAACCTGAAGTC
E L H I F G S I N G V D F D M V G Q G T G N P N D G Y E E L N L K S

mNeonGreen

CACCAAGGGTGACCTCCAGTTCTCCCCCTGGATTCTGGTCCCTCATATCGGGTATGGCTTCCATCAGTACCTGCCCTACCCTGACGGGATGTCGCCTTTC
T K G D L Q F S P W I L V P H I G Y G F H Q Y L P Y P D G M S P F

mNeonGreen

CAGGCCGCCATGGTAGATGGCTCCGGATACCAAGTCCATCGCACAATGCAGTTTGAAGATGGTGCCTCCCTTACTGTTAACTACCGCTACACCTACGAGG
Q A A M V D G S G Y Q V H R T M Q F E D G A S L T V N Y R Y T Y E

mNeonGreen

GAAGCCACATCAAAGGAGAGGCCCAGGTGAAGGGGACTGGTTTCCCTGCTGACGGTCCTGTGATGACCAACTCGCTGACCGCTGCGGACTGGTGCAGGTC
G S H I K G E A Q V K G T G F P A D G P V M T N S L T A A D W C R S

mNeonGreen

GAAGAAGACTTACCCCAACGACAAAACCATCATCAGTACCTTTAAGTGGAGTTACACCACTGGAAATGGCAAGCGCTACAGGAGCACTGCGCGGACCACC
K K T Y P N D K T I I S T F K W S Y T T G N G K R Y R S T A R T T

mNeonGreen

TACACCTTTGCCAAGCCAATGGCGGCTAACTATCTGAAGAACCAGCCGATGTACGTGTTCCGTAAGACGGAGCTCAAGCACTCCAAGACCGAGCTCAACT
Y T F A K P M A A N Y L K N Q P M Y V F R K T E L K H S K T E L N

mNeonGreen | Kir2.1 membrane trafficking sequence

TCAAGGAGTGGCAAAAGGCCTTTACCGATGTGATGGGCATGGACGAGCTGTACAAGAAGAGCAGGATCACCAGCGAGGGCGAGTACATCCCCCTGGACCA
F K E W Q K A F T D V M G M D E L Y K K S R I T S E G E Y I P L D Q

Kir2.1 ER export sequence | Kv2.1 proximal restriction sequence

GATCGACATCAACGTGTTCTGCTACGAGAACGAGGTGCAAAGTCAGCCTATCCTGAACACAAAGGAAATGGCTCCACAGTCTAAGCCTCCCGAAGAGCTT
I D I N V F C Y E N E V Q S Q P I L N T K E M A P Q S K P P E E L

Kv2.1 proximal restriction sequence

GAGATGTCCAGTATGCCAAGTCCCGTGGCTCCCCTCCCTGCCAGGACTGAAGGAGTGATTGACATGAGGAGTATGTCATCTATTGATAGCTTCATCTCTT
E M S S M P S P V A P L P A R T E G V I D M R S M S S I D S F I S

Kv2.1 proximal restriction sequence

GCGCAACAGATTTCCCCGAGGCTACTCGATTCTAA
C A T D F P E A T R F .

**Figure S3 |** DNA and amino acid sequences of Ace-mNeon2 fused to a somatic restriction sequence from Kv2.1.

Ace opsin

ATGGCTGACGTGGAAACCGAGACCGGCATGATTGCACAGTGGATTGTCTTTGCTATTATGGCTGCTGCTGCTATTGCTTTTGGAGTGGCTGTGCACTTTC
M A D V E T E T G M I A Q W I V F A I M A A A A I A F G V A V H F

Ace opsin

GGCCTTCAGAGCTGAAGAGCGCATACTATATCAACATTGCCATCTGCACTATCGCCGCTACCGCTTACTATGCAATGGCCGTGAACTACCAGGACCTGAC
R P S E L K S A Y Y I N I A I C T I A A T A Y Y A M A V N Y Q D L T

Ace opsin

AATGAATGGTGAAAGGCAGGTGGTCTACGCAAGATATATTAGCTGGGTGCTGACCACACCACTGCTCCTGCTCGATCTCATCGTCATGACCAAGATGGGC
M N G E R Q V V Y A R Y I S W V L T T P L L L L D L I V M T K M G

Ace opsin

GGAGTGATGATTTCTTGGGTCATCGGCGCAGACATTTTCATGATCGTGTTTGGTATTCTGGGCGCCTTCGAGGATGAACACAAGTTCAAATGGGTGTACT
G V M I S W V I G A D I F M I V F G I L G A F E D E H K F K W V Y

Ace opsin

TTATCGCTGGATGTGTGATGCAGGCAGTCCTGACATACGGGATGTATAACGCCACTTGGAAAGACGATCTGAAGAAAAGCCCCGAGTACCATAGCTCCTA
F I A G C V M Q A V L T Y G M Y N A T W K D D L K K S P E Y H S S Y

Ace opsin

TGTCAGTCTGCTCGTCTTCCTGTCAATCCTCTGGGTGTTTTATCCTGTCGTGTGGGCTTTCGGGTCTGGTAGTGGCGTGCTGTCCGTCGACAATGAGGCC
V S L L V F L S I L W V F Y P V V W A F G S G S G V L S V D N E A

Ace opsin | Linker | mRuby3

ATTCTCATGGGAATCCTGGATGTGCTCGCTAAGCCACTGTTTGGAATGGGGTGCCTCATTGCCCATGAGACTATCTTCAAGTCCATCCTGATTAAGGAAA
I L M G I L D V L A K P L F G M G C L I A H E T I F K S I L I K E

mRuby3

ATATGCGGATGAAGGTCGTGATGGAAGGGTCTGTCAATGGGCACCAGTTCAAGTGCACCGGAGAGGGAGAGGGCAGGCCATACGAGGGCGTGCAGACAAT
N M R M K V V M E G S V N G H Q F K C T G E G E G R P Y E G V Q T M

mRuby3

GAGGATCAAAGTGATCGAGGGAGGACCACTGCCTTTCGCCTTTGACATCCTGGCCACCAGCTTCATGTACGGCAGCAGGACCTTCATCAAGTATCCAGCC
R I K V I E G G P L P F A F D I L A T S F M Y G S R T F I K Y P A

mRuby3

GACATCCCCGATTTCTTTAAGCAGAGCTTCCCCGAGGGCTTTACCTGGGAGAGGGTGACAAGATACGAGGATGGCGGCGTGGTGACCGTGACACAGGACA
D I P D F F K Q S F P E G F T W E R V T R Y E D G G V V T V T Q D

mRuby3

CCTCTCTGGAGGATGGCGAGCTGGTGTATAACGTGAAGGTGAGGGGCGTGAACTTCCCTAGCAATGGCCCAGTGATGCAGAAGAAGACCAAGGGCTGGGA
T S L E D G E L V Y N V K V R G V N F P S N G P V M Q K K T K G W E

mRuby3

GCCCAATACAGAGATGATGTACCCTGCAGACGGAGGCCTGAGGGGATATACCGACATCGCCCTGAAGGTGGATGGAGGAGGACACCTGCACTGTAACTTT
P N T E M M Y P A D G G L R G Y T D I A L K V D G G G H L H C N F

mRuby3

GTGACCACATACCGCTCCAAGAAGACAGTGGGCAATATCAAGATGCCAGGAGTGCACGCCGTGGACCACAGGCTGGAGCGCATCGAGGAGTCTGATAACG
V T T Y R S K K T V G N I K M P G V H A V D H R L E R I E E S D N

mRuby3

AGACATATGTGGTGCAGAGAGAGGTGGCCGTGGCCAAGTACTCTAATCTGGGCGGCGGGATGGACGAGCTGTATAAGAAGAGCAGGATCACCAGCGAGGG
E T Y V V Q R E V A V A K Y S N L G G G M D E L Y K K S R I T S E G

Kir2.1 membrane trafficking sequence | Kir2.1 ER export sequence | Kv2.1 proximal restriction sequence

CGAGTACATCCCCCTGGACCAGATCGACATCAACGTGTTCTGCTACGAGAACGAGGTGCAAAGTCAGCCTATCCTGAACACAAAGGAAATGGCTCCACAG
E Y I P L D Q I D I N V F C Y E N E V Q S Q P I L N T K E M A P Q

Kv2.1 proximal restriction sequence

TCTAAGCCTCCCGAAGAGCTTGAGATGTCCAGTATGCCAAGTCCCGTGGCTCCCCTCCCTGCCAGGACTGAAGGAGTGATTGACATGAGGAGTATGTCAT
S K P P E E L E M S S M P S P V A P L P A R T E G V I D M R S M S

Kv2.1 proximal restriction sequence

CTATTGATAGCTTCATCTCTTGCGCAACAGATTTCCCCGAGGCTACTCGATTCTAA
S I D S F I S C A T D F P E A T R F .

**Figure S4 |** DNA and amino acid sequences of VARNAM2 fused to a somatic restriction sequence from Kv2.1.

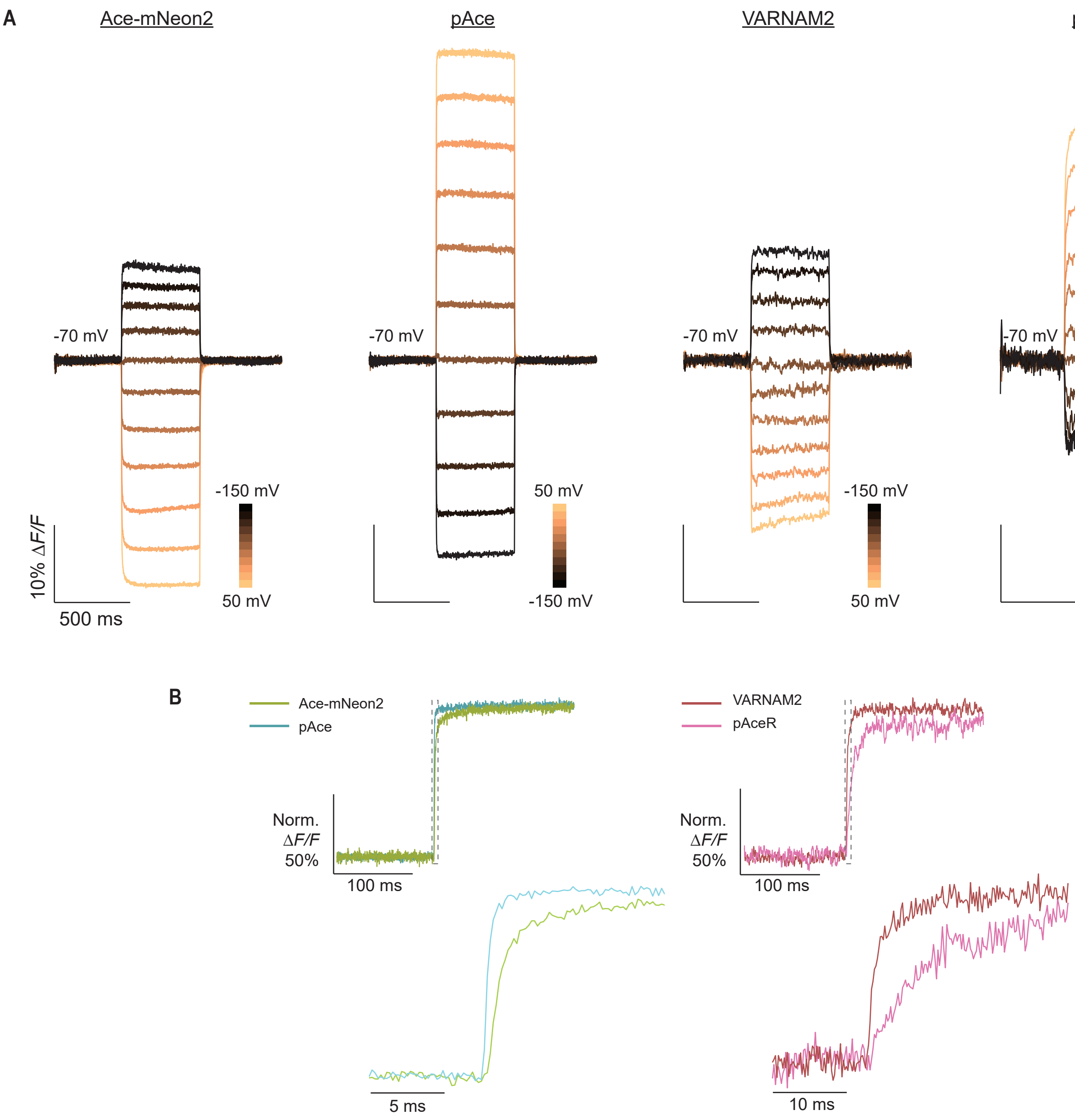


**Figure S5 |** Voltage sensitivity and kinetics of the four FRET-opsin indicators in HEK cells. (A) Example fluorescence responses from a HEK cell expressing Ace-mNeon2, pAce, VARNAM2 or pAceR during whole-cell voltage clamp recordings. HEK cells were held at -70 mV responses were recorded to depolarizing and hyperpolarizing voltage steps at 20 mV increments. (B) Normalized fluorescence responses of HEK cells transfected with Ace-mNeon2 or pAce (*left*), and VARNAM2 or pAceR (*right*) during 120 mV depolarization. Dashed grey box represents interval shown below at an expanded time scale. Imaging conditions: 505 nm LED for Ace-mNeon2 and pAce and 565 nm LED for VARNAM2 and pAceR, 25 mW mm$^{-2}$ at sample plane; image acquisition: 1 kHz.

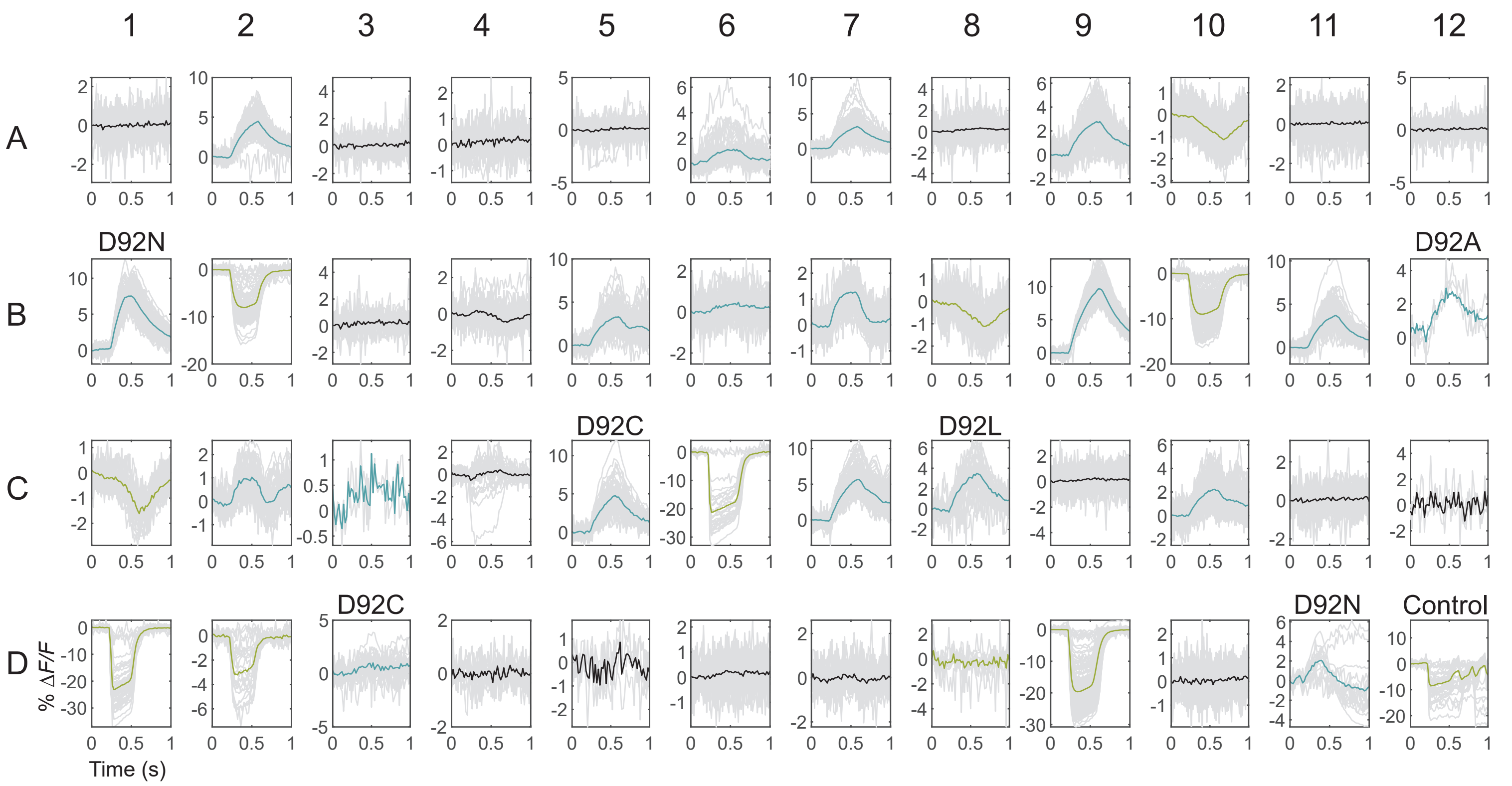


**Figure S6 |** High-throughput voltage screening of Ace-mNeon2 D92X mutants. 48 variants containing saturation mutations at Ace D92 were each transfected in electrically excitable HEK cells[25,26] and screened for voltage sensitivity on the high-throughput platform[18]. Shown here are representative fluorescence responses of all neurons (grey) from one field-of-view and one round of screening per variant. The average responses to depolarizing field potentials are indicated in blue for positive-polarity, green for negative-polarity and black for non-responding variants. Mutational information is indicated on top of each well for the positive-polarity variants, which were chosen for sequencing post-screening. (n>100 cells/variant).

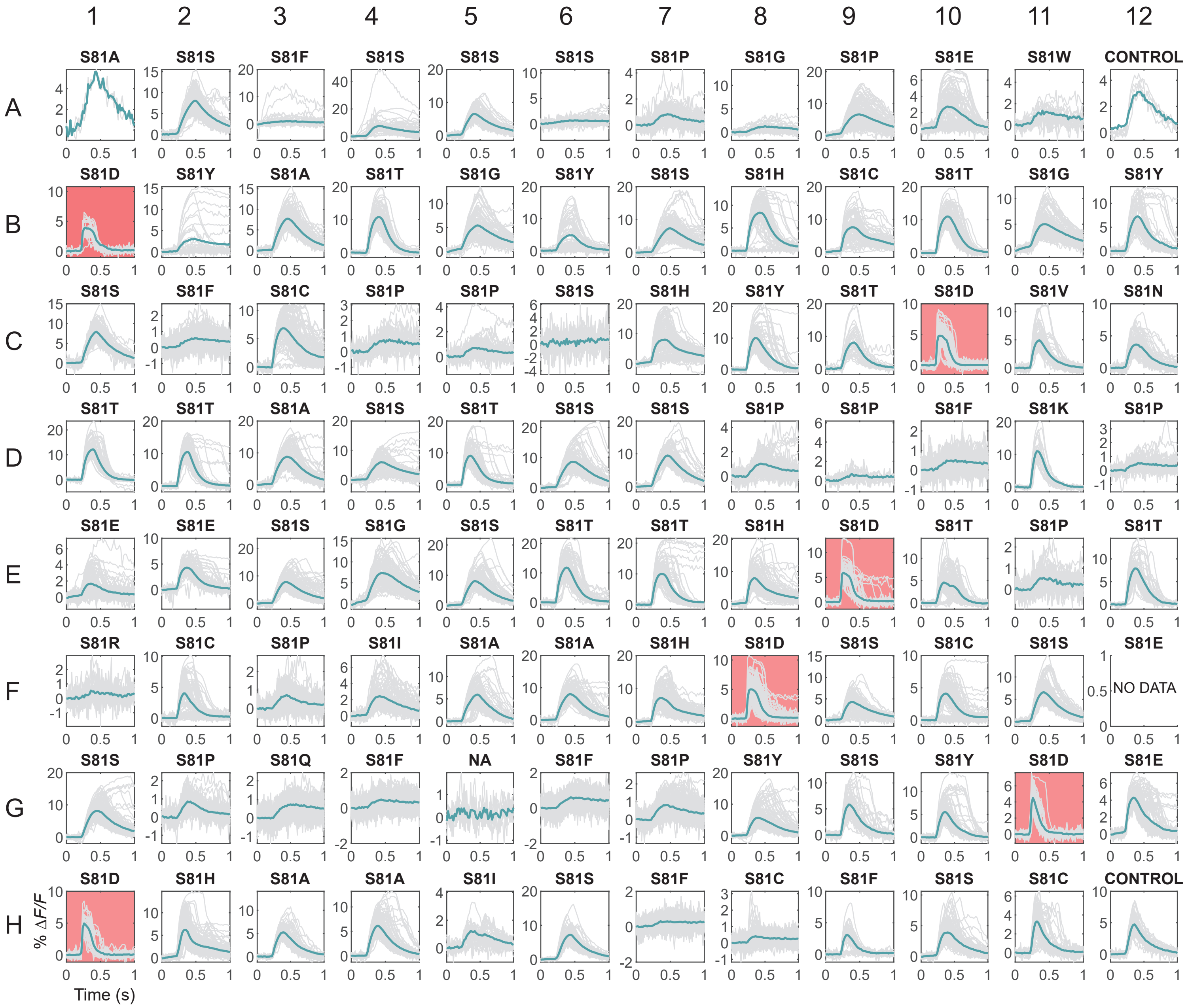


**Figure S7** | High-throughput screening for kinetics rescue mutations in Ace-mNeon2 D92N. 96 variants containing saturation mutations at Ace S81 were each transfected in electrically excitable HEK cells[25,26] and screened for voltage sensitivity and response kinetics on the high-throughput platform[18]. Shown here are representative fluorescence responses of all neurons (grey) from one field-of-view and one round of screening per variant. The average responses to depolarizing field potentials are indicated in blue. Mutational information is indicated on top of each well (n>100 cells/variant except A1 and A12, where n<10 cells/well). Wells shaded in pink exhibited improvements in response kinetics, which is more conspicuous in the maximum response races. Images were acquired at 200 Hz for this experiment (see Methods).

**A** Ace R78X and W178X on Ace-mNeon2 S81D D92N

| | 1 | 2 | 3 | 4 | 5 | 6 | 7 | 8 | 9 | 10 | 11 | 12 |
|---|---|---|---|---|---|---|---|---|---|---|---|---|
| **A** | 6<br>R78V | 6<br>R78A | 8<br>R78F | 27<br>R78X | 9<br>R78V | 7<br>R78L | 27<br>R78C | 3<br>R78W | 7<br>R78Q | 6<br>R78S | 1<br>R78F | Control |
| **B** | 6<br>R78H | 13<br>R78M | 8<br>R78G | 8<br>R78W | 6<br>R78Y | 2<br>R78Y | 3<br>R78Y | 10<br>R78M | 37<br>R78K | 43<br>R78K | 6<br>R78V | 7<br>R78L |
| **C** | 8<br>R78R | 8<br>R78W | 8<br>R78S | 10<br>R78R | 0<br>R78W | 35<br>R78K | 2<br>R78W | 3<br>R78A | 3<br>R78C | 2<br>R78Y | 4<br>R78L | 5<br>R78V |
| **D** | 1<br>R78W | 8<br>R78L | 8<br>R78G | 0<br>R78D | 8<br>R78I | 2<br>R78Y | 2<br>R78W | 7<br>R78R | 2<br>R78C | 4<br>R78Y | 3<br>R78C | 15<br>R78N |
| **E** | 5<br>W178A | 10<br>W178 | 8<br>W178W | 5<br>W178A | 4<br>W178K | 2<br>No data | 10<br>W178W | 4<br>W178M | 6<br>W178V | 8<br>W178G | 11<br>W178F | 5<br>W178M |
| **F** | 7 | 6<br>W178Y | 6<br>W178S | 5<br>W178A | 8 | 11<br>W178V | 10<br>W178F | 11<br>W178H | 5<br>W178M | 8<br>W178W | 8<br>W178W | 21<br>W178Q |
| **G** | 7<br>W178Y | 4<br>W178I | 9<br>W178S | 6<br>W178V | 10<br>W178W | 11<br>W178S | 21<br>W178E | 0<br>W178P | 14<br>W178S | 12<br>W178G | 12<br>W178L | 4<br>W178I |
| **H** | 19<br>W178N | 13<br>W178Q | 7<br>W178S | 9<br>W178C | 10<br>W178W | 5<br>W178I | 6<br>W178F | 0<br>No data | 5<br>W178T | 8<br>W178V | 10<br>W178H | Control |

**B** Ace W178X on Ace-mNeon2 S81D D92N R78K (well B10 from above)

| | 1 | 2 | 3 | 4 | 5 | 6 | 7 | 8 | 9 | 10 | 11 | 12 |
|---|---|---|---|---|---|---|---|---|---|---|---|---|
| **A** | 40<br>W178Y | 11<br>W178Q | 8<br>W178N | 15<br>W178C | 40<br>W178L | 38<br>W178W | 8<br>W178S | 40<br>W178W | 0<br>No data | 31<br>W178C | 32<br>W178V | 16<br>W178N |
| **B** | 39<br>W178Y | 27<br>W178K | 18<br>W178N | 39<br>W178Y | 35<br>W178Y | 24<br>W178K | 32<br>W178I | 23<br>W178G | 40<br>W178Y | 28<br>W178T | 42<br>W178W | 25<br>W178C |
| **C** | 0<br>W178R | 0<br>W178P | 25<br>W178H | 35<br>W178V | 46<br>W178F | 25<br>W178G | 30<br>W178M | 42<br>W178W | 36<br>W178Y | 41<br>W178W | 37<br>W178W | 25<br>W178C |
| **D** | 40<br>W178W | 26<br>W178I | 43<br>W178T | 38<br>W178W | 45<br>W178F | 12<br>W178D | 21<br>W178P | 34<br>W178V | 31<br>W178T | 40<br>W178V | 46<br>W178C | Control |

**Figure S8 |** High-throughput screening for voltage sensitivity of positive-polarity variant Ace-mNeon2 S81D and D92N.
(A) 48 variants containing saturation mutations at Ace R78 or W178 on the backbone of Ace-mNeon2 S81D D92N were each transfected in electrically excitable HEK cells[25,26] and screened for voltage sensitivity on the high-throughput platform[18]. The maximum %$\Delta F/F$ obtained for each variant across 4 independent rounds of screening are indicated in each well together with the sequence information. Color density corresponds to the size of the signal in the positive direction.
(B) Same as above for Ace W178X mutagenesis on Ace-mNeon2 S81D D92N R78K backbone.

Ace opsin

ATGGCTGACGTGGAAACCGAGACCGGCATGATTGCACAGTGGATTGTCTTTGCTATTATGGCTGCTGCTGCTATTGCTTTTGGAGTGGCTGTGCACTTTC
M A D V E T E T G M I A Q W I V F A I M A A A A I A F G V A V H F

Ace opsin

GGCCTTCAGAGCTGAAGAGCGCATACTATATCAACATTGCCATCTGCACTATCGCCGCTACCGCTTACTATGCAATGGCCGTGAACTACCAGGACCTGAC
R P S E L K S A Y Y I N I A I C T I A A T A Y Y A M A V N Y Q D L T

Ace opsin

AATGAATGGTGAAAGGCAGGTGGTCTACGCAAAGTATATTGACTGGGTGCTGACCACACCACTGCTCCTGCTCAACCTCATCGTCATGACCAAGATGGGC
M N G E R Q V V Y A K Y I D W V L T T P L L L L N L I V M T K M G

Ace opsin

GGAGTGATGATTTCTTGGGTCATCGGCGCAGACATTTTCATGATCGTGTTTGGTATTCTGGGCGCCTTCGAGGATGAACACAAGTTCAAATGGGTGTACT
G V M I S W V I G A D I F M I V F G I L G A F E D E H K F K W V Y

Ace opsin

TTATCGCTGGATGTGTGATGCAGGCAGTCCTGACATACGGGATGTATAACGCCACTTGGAAAGACGATCTGAAGAAAAGCCCCGAGTACCATAGCTCCTA
F I A G C V M Q A V L T Y G M Y N A T W K D D L K K S P E Y H S S Y

Ace opsin

TGTCAGTCTGCTCGTCTTCCTGTCAATCCTCTTCGTGTTTTATCCTGTCGTGTGGGCTTTCGGGTCTGGTAGTGGCGTGCTGTCCGTCGACAATGAGGCC
V S L L V F L S I L F V F Y P V V W A F G S G S G V L S V D N E A

Ace opsin | Linker | mNeonGreen

ATTCTCATGGGAATCCTGGATGTGCTCGCTAAGCCACTGTTTGGAATGGGGTGCCTCATTGCCCATGAGACTATCTTCAAGTCCTACCCAGCGACACATG
I L M G I L D V L A K P L F G M G C L I A H E T I F K S Y P A T H

mNeonGreen

AGTTACACATCTTTGGCTCCATCAACGGTGTGGACTTTGACATGGTGGGTCAGGGCACCGGCAATCCAAATGATGGTTATGAGGAGTTAAACCTGAAGTC
E L H I F G S I N G V D F D M V G Q G T G N P N D G Y E E L N L K S

mNeonGreen

CACCAAGGGTGACCTCCAGTTCTCCCCCTGGATTCTGGTCCCTCATATCGGGTATGGCTTCCATCAGTACCTGCCCTACCCTGACGGGATGTCGCCTTTC
T K G D L Q F S P W I L V P H I G Y G F H Q Y L P Y P D G M S P F

mNeonGreen

CAGGCCGCCATGGTAGATGGCTCCGGATACCAAGTCCATCGCACAATGCAGTTTGAAGATGGTGCCTCCCTTACTGTTAACTACCGCTACACCTACGAGG
Q A A M V D G S G Y Q V H R T M Q F E D G A S L T V N Y R Y T Y E

mNeonGreen

GAAGCCACATCAAAGGAGAGGCCCAGGTGAAGGGGACTGGTTTCCCTGCTGACGGTCCTGTGATGACCAACTCGCTGACCGCTGCGGACTGGTGCAGGTC
G S H I K G E A Q V K G T G F P A D G P V M T N S L T A A D W C R S

mNeonGreen

GAAGAAGACTTACCCCAACGACAAAACCATCATCAGTACCTTTAAGTGGAGTTACACCACTGGAAATGGCAAGCGCTACAGGAGCACTGCGCGGACCACC
K K T Y P N D K T I I S T F K W S Y T T G N G K R Y R S T A R T T

mNeonGreen

TACACCTTTGCCAAGCCAATGGCGGCTAACTATCTGAAGAACCAGCCGATGTACGTGTTCCGTAAGACGGAGCTCAAGCACTCCAAGACCGAGCTCAACT
Y T F A K P M A A N Y L K N Q P M Y V F R K T E L K H S K T E L N

mNeonGreen | Kir2.1 membrane trafficking sequence

TCAAGGAGTGGCAAAAGGCCTTTACCGATGTGATGGGCATGGACGAGCTGTACAAGAAGAGCAGGATCACCAGCGAGGGCGAGTACATCCCCCTGGACCA
F K E W Q K A F T D V M G M D E L Y K K S R I T S E G E Y I P L D Q

Kir2.1 ER export sequence | Kv2.1 proximal restriction sequence

GATCGACATCAACGTGTTCTGCTACGAGAACGAGGTGCAAAGTCAGCCTATCCTGAACACAAAGGAAATGGCTCCACAGTCTAAGCCTCCCGAAGAGCTT
I D I N V F C Y E N E V Q S Q P I L N T K E M A P Q S K P P E E L

Kv2.1 proximal restriction sequence

GAGATGTCCAGTATGCCAAGTCCCGTGGCTCCCCTCCCTGCCAGGACTGAAGGAGTGATTGACATGAGGAGTATGTCATCTATTGATAGCTTCATCTCTT
E M S S M P S P V A P L P A R T E G V I D M R S M S S I D S F I S

Kv2.1 proximal restriction sequence

GCGCAACAGATTTCCCCGAGGCTACTCGATTCTAA
C A T D F P E A T R F .

**Figure S9** | DNA and amino acid sequences of PACE fused to a somatic restriction sequence from Kv2.1.

Ace opsin

ATGGCTGACGTGGAAACCGAGACCGGCATGATTGCACAGTGGATTGTCTTTGCTATTATGGCTGCTGCTGCTATTGCTTTTGGAGTGGCTGTGCACTTTC
M A D V E T E T G M I A Q W I V F A I M A A A A I A F G V A V H F

Ace opsin

GGCCTTCAGAGCTGAAGAGCGCATACTATATCAACATTGCCATCTGCACTATCGCCGCTACCGCTTACTATGCAATGGCCGTGAACTACCAGGACCTGAC
R P S E L K S A Y Y I N I A I C T I A A T A Y Y A M A V N Y Q D L T

Ace opsin

AATGAATGGTGAAAGGCAGGTGGTCTACGCAGAGTATATTGACTGGGTGCTGACCACACCACTGCTCCTGCTCAACCTCATCGTCATGACCAAGATGGGC
M N G E R Q V V Y A E Y I D W V L T T P L L L L N L I V M T K M G

Ace opsin

GGAGTGATGATTTCTTGGGTCATCGGCGCAGACATTTTCATGATCGTGTTTGGTATTCTGGGCGCCTTCGAGGATGAACACAAGTTCAAATGGGTGTACT
G V M I S W V I G A D I F M I V F G I L G A F E D E H K F K W V Y

Ace opsin

TTATCGCTGGATGTGTGATGCAGGCAGTCCTGACATACGGGATGTATAACGCCACTTGGAAAGACGATCTGAAGAAAAGCCCCGAGTACCATAGCTCCTA
F I A G C V M Q A V L T Y G M Y N A T W K D D L K K S P E Y H S S Y

Ace opsin

TGTCAGTCTGCTCGTCTTCCTGTCAATCCTCTGGGTGTTTTATCCTGTCGTGTGGGCTTTCGGGTCTGGTAGTGGCGTGCTGTCCGTCGACAATGAGGCC
V S L L V F L S I L W V F Y P V V W A F G S G S G V L S V D N E A

Ace opsin | Linker | mRuby3

ATTCTCATGGGAATCCTGGATGTGCTCGCTAAGCCACTGTTTGGAATGGGGTGCCTCATTGCCCATGAGACTATCTTCAAGTCCATCCTGATTAAGGAAA
I L M G I L D V L A K P L F G M G C L I A H E T I F K S I L I K E

mRuby3

ATATGCGGATGAAGGTCGTGATGGAAGGGTCTGTCAATGGGCACCAGTTCAAGTGCACCGGAGAGGGAGAGGGCAGGCCATACGAGGGCGTGCAGACAAT
N M R M K V V M E G S V N G H Q F K C T G E G E G R P Y E G V Q T M

mRuby3

GAGGATCAAAGTGATCGAGGGAGGACCACTGCCTTTCGCCTTTGACATCCTGGCCACCAGCTTCATGTACGGCAGCAGGACCTTCATCAAGTATCCAGCC
R I K V I E G G P L P F A F D I L A T S F M Y G S R T F I K Y P A

mRuby3

GACATCCCCGATTTCTTTAAGCAGAGCTTCCCCGAGGGCTTTACCTGGGAGAGGGTGACAAGATACGAGGATGGCGGCGTGGTGACCGTGACACAGGACA
D I P D F F K Q S F P E G F T W E R V T R Y E D G G V V T V T Q D

mRuby3

CCTCTCTGGAGGATGGCGAGCTGGTGTATAACGTGAAGGTGAGGGGCGTGAACTTCCCTAGCAATGGCCCAGTGATGCAGAAGAAGACCAAGGGCTGGGA
T S L E D G E L V Y N V K V R G V N F P S N G P V M Q K K T K G W E

mRuby3

GCCCAATACAGAGATGATGTACCCTGCAGACGGAGGCCTGAGGGGATATACCGACATCGCCCTGAAGGTGGATGGAGGAGGACACCTGCACTGTAACTTT
P N T E M M Y P A D G G L R G Y T D I A L K V D G G G H L H C N F

mRuby3

GTGACCACATACCGCTCCAAGAAGACAGTGGGCAATATCAAGATGCCAGGAGTGCACGCCGTGGACCACAGGCTGGAGCGCATCGAGGAGTCTGATAACG
V T T Y R S K K T V G N I K M P G V H A V D H R L E R I E E S D N

mRuby3

AGACATATGTGGTGCAGAGAGAGGTGGCCGTGGCCAAGTACTCTAATCTGGGCGGCGGGATGGACGAGCTGTATAAGAAGAGCAGGATCACCAGCGAGGG
E T Y V V Q R E V A V A K Y S N L G G G M D E L Y K K S R I T S E G

Kir2.1 membrane trafficking sequence | Kir2.1 ER export sequence | Kv2.1 proximal restriction sequence

CGAGTACATCCCCCTGGACCAGATCGACATCAACGTGTTCTGCTACGAGAACGAGGTGCAAAGTCAGCCTATCCTGAACACAAAGGAAATGGCTCCACAG
E Y I P L D Q I D I N V F C Y E N E V Q S Q P I L N T K E M A P Q

Kv2.1 proximal restriction sequence

TCTAAGCCTCCCGAAGAGCTTGAGATGTCCAGTATGCCAAGTCCCGTGGCTCCCCTCCCTGCCAGGACTGAAGGAGTGATTGACATGAGGAGTATGTCAT
S K P P E E L E M S S M P S P V A P L P A R T E G V I D M R S M S

Kv2.1 proximal restriction sequence

CTATTGATAGCTTCATCTCTTGCGCAACAGATTTCCCCGAGGCTACTCGATTCTAA
S I D S F I S C A T D F P E A T R F .

**Figure S10 |** DNA and amino acid sequences of PACER fused to a somatic restriction sequence from Kv2.1.

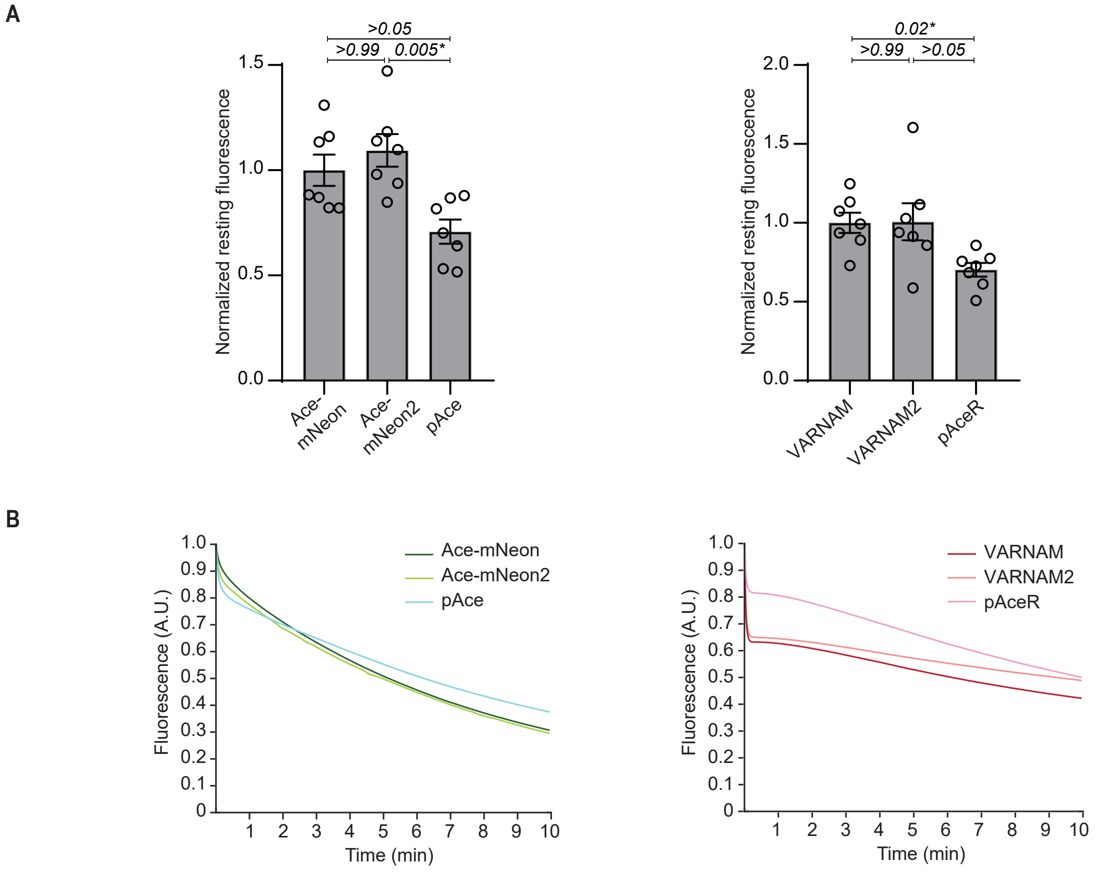


**Figure S11 |** Brightness and photobleaching characteristics of the FRET-opsin indicators.
(A) *Left*, Resting fluorescence of Ace-mNeon, Ace-mNeon2 and pAce normalized to average resting intensity of Ace-mNeon. Values represent mean ± S.E.M. P values are italicized. Statistical comparisons were made across all conditions. Asterisks denote significance (Kruskal-Wallis test with Dunn's multiple comparisons correction). *Right*, same as above for VARNAM, VARNAM2 and pAceR (n=7 wells/condition, ~500 cells/well).
(B) Photobleaching profiles of Ace-mNeon, Ace-mNeon2 and pAce (*left*) and VARNAM, VARNAM2 and pAceR (*right*) in HEK cells under continuous illumination (n=4 wells each, ~100 cells/well) Imaging conditions: 505 nm LED for the green indicators and 565 nm LED for the red indicators, 25 mW mm$^{-2}$ at sample plane.

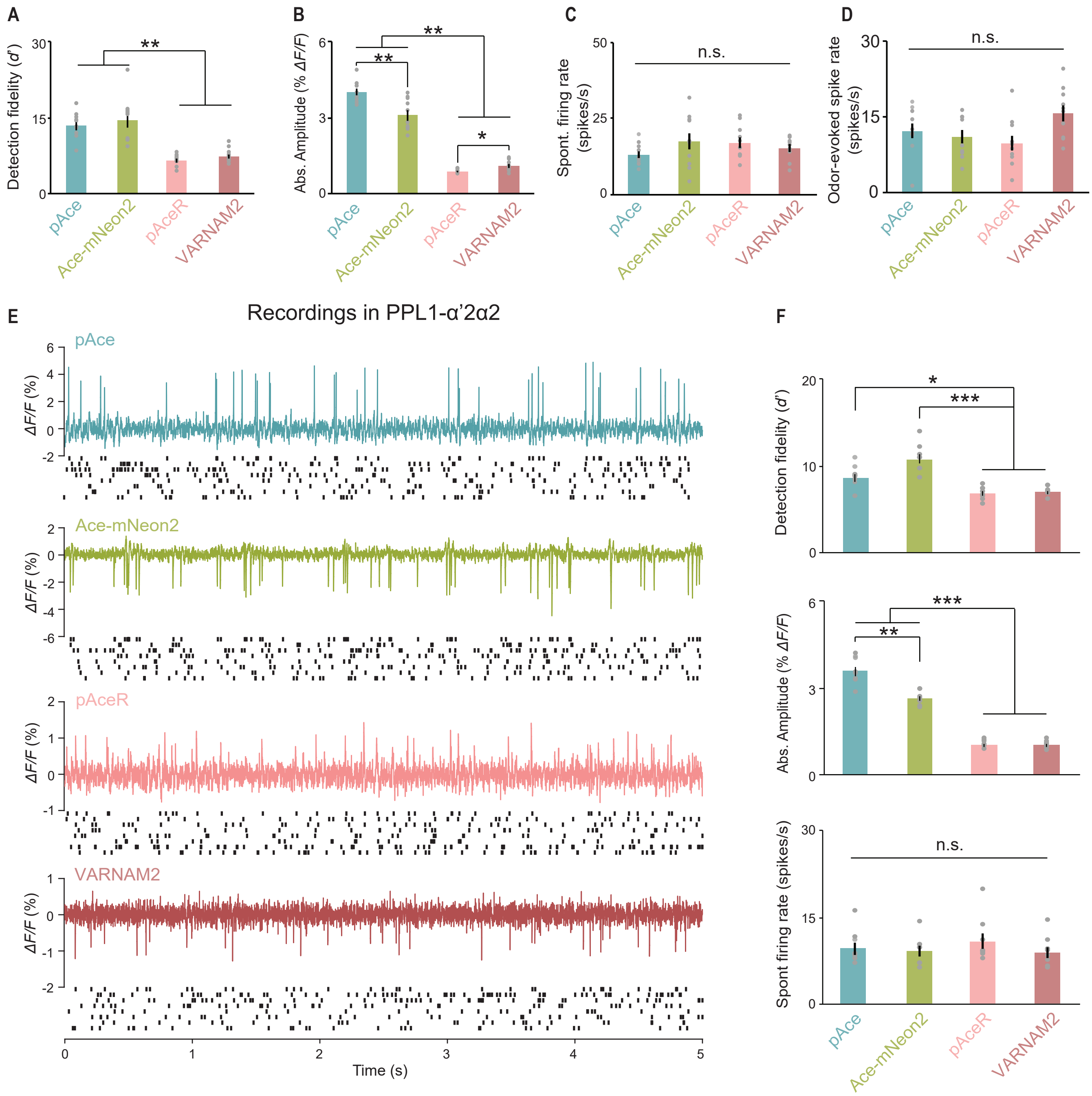


**Figure S12** | Characterization of the FRET-opsin indicators in PPL1-γ2α'1 and PPL1-α'2α2 neurons in *Drosophila* . (A-D) Comparisons of (A) spike detection fidelities (B) absolute amplitudes (C) mean spontaneous firing rates, and (D) odor-evoked firing rate change obtained from recordings in PPL1-γ2α'1 expressing each of the four indicators. (n = 10 trials; 2 trials per fly; **P*<0.05, ***P*<0.01, ****P*<0.001, n.s.=not significant, Kruskal-Wallis ANOVA and post-hoc Mann-Whitney U-tests with Holm-Bonferroni correction).
(E) Representative optical recordings (*top*) and raster plots (*bottom*) of 5-s spontaneous spiking in a PPL1-α'2α2 neuron expressing pAce, Ace-mNeon2, pAceR or VARNAM2 (n = 8 trials; 2 trials per fly).
(F) Comparisons of detection fidelities (*top*), absolute amplitudes (*center*), and mean spontaneous firing rates (*bottom*) in PPL1-α'2α2 neurons. (n = 8 trials; 2 trials per fly; **P*<0.05, ***P*<0.01, ****P*<0.001, Kruskal-Wallis ANOVA and post-hoc Mann-Whitney U-tests with Holm-Bonferroni correction).

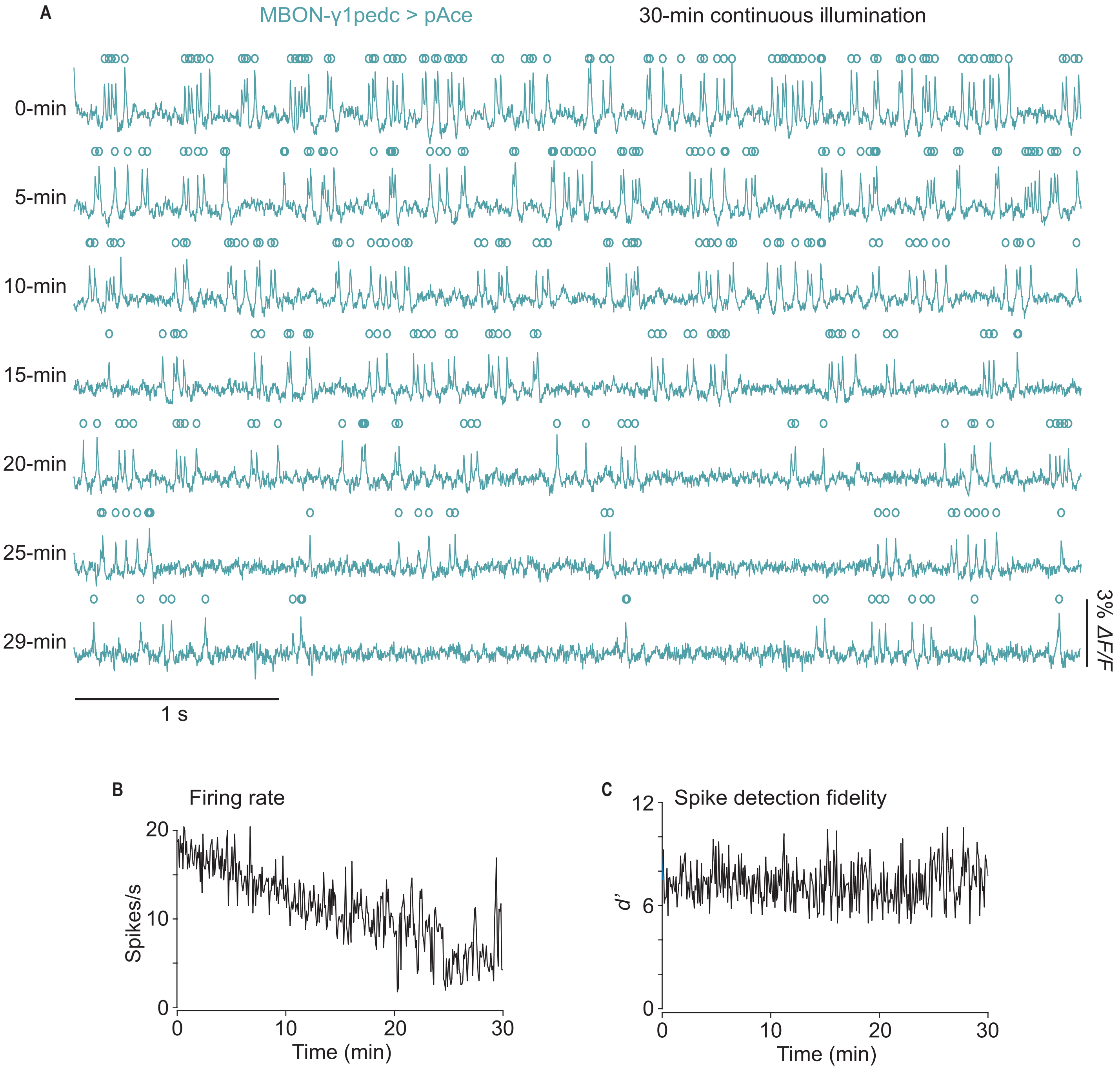


**Figure S13 |** 30-min continuous imaging using pAce in *Drosophila*.
(A) Example 5 s recordings collected at regular time intervals during 30-min continuous illumination showing spontaneous spiking in a MBON-γ1pedc>α/β neuron expressing pAce.
(B) Time-varying spike rate in the recorded MBON-γ1pedc>α/β neuron during the 30-min imaging session.
(C) Spike detection fidelity in the recorded MBON-γ1pedc>α/β neuron during the 30-min imaging session.

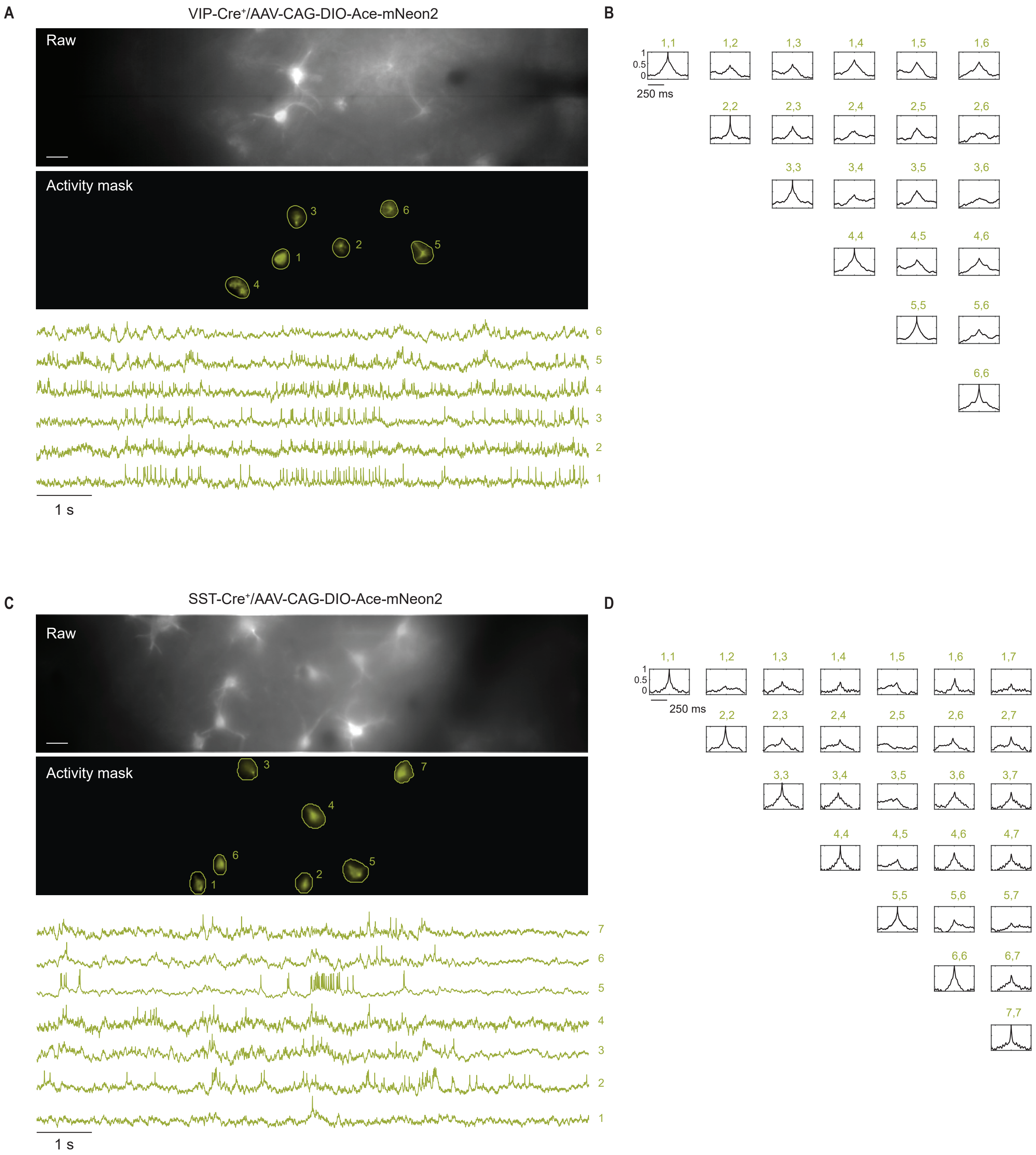
A
VIP-Cre+/AAV-CAG-DIO-Ace-mNeon2
Raw
Activity mask
1 2 3 4 5 6
1 s
B
1,1 1,2 1,3 1,4 1,5 1,6
2,2 2,3 2,4 2,5 2,6
3,3 3,4 3,5 3,6
4,4 4,5 4,6
5,5 5,6
6,6
1
0.5
0
250 ms
C
SST-Cre+/AAV-CAG-DIO-Ace-mNeon2
Raw
Activity mask
1 2 3 4 5 6 7
1 s
D
1,1 1,2 1,3 1,4 1,5 1,6 1,7
2,2 2,3 2,4 2,5 2,6 2,7
3,3 3,4 3,5 3,6 3,7
4,4 4,5 4,6 4,7
5,5 5,6 5,7
6,6 6,7
7,7
1
0.5
0
250 ms

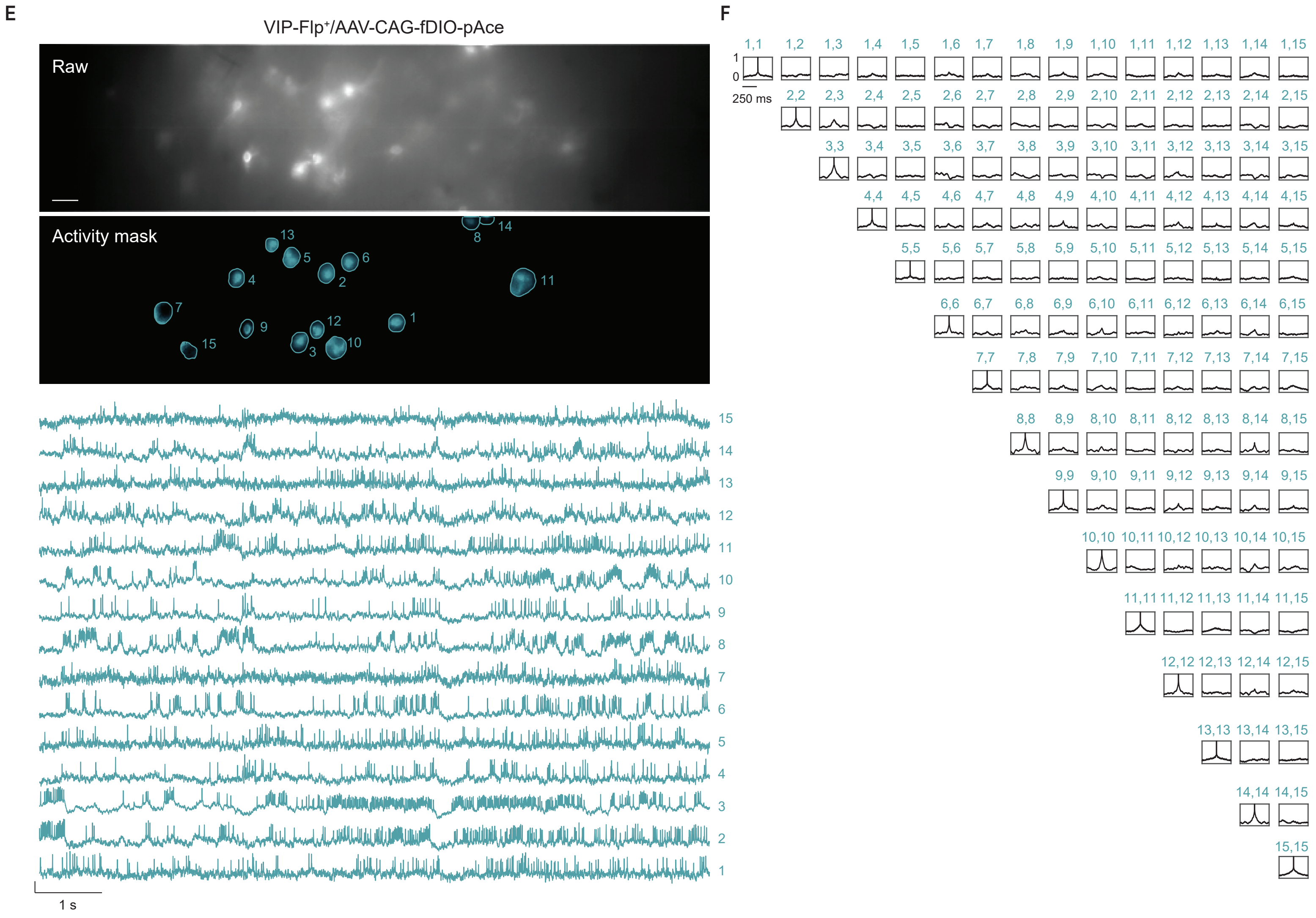


**Figure S14 |** Minimal cross-contamination of voltage signals from adjacent cells in widefield recordings.
(A) *Top*, Representative epifluorescence image of a single field-of-view from a VIP-Cre+ mouse, expressing *Cre*-dependent soma-targeted Ace-mNeon2 in V1. Scale bar: 50 µm. *Center*, mask image showing active ROIs. *Bottom*, *ΔF/F* traces showing spontaneous activity from the ROIs numbered in the mask image.
(B) Auto- and cross-correlations of the *ΔF/F* traces for neuronal pairs in the FoV above.
(C) Same as (A) for SST-Cre+ mouse.
(D) Same as (B) for the FoV in (C).
(E) Same as (A) for VIP-Flp+ mouse expressing *Flp*-dependent soma-targeted pAce in V1.
(F) Same as (B) for the FoV in (E).

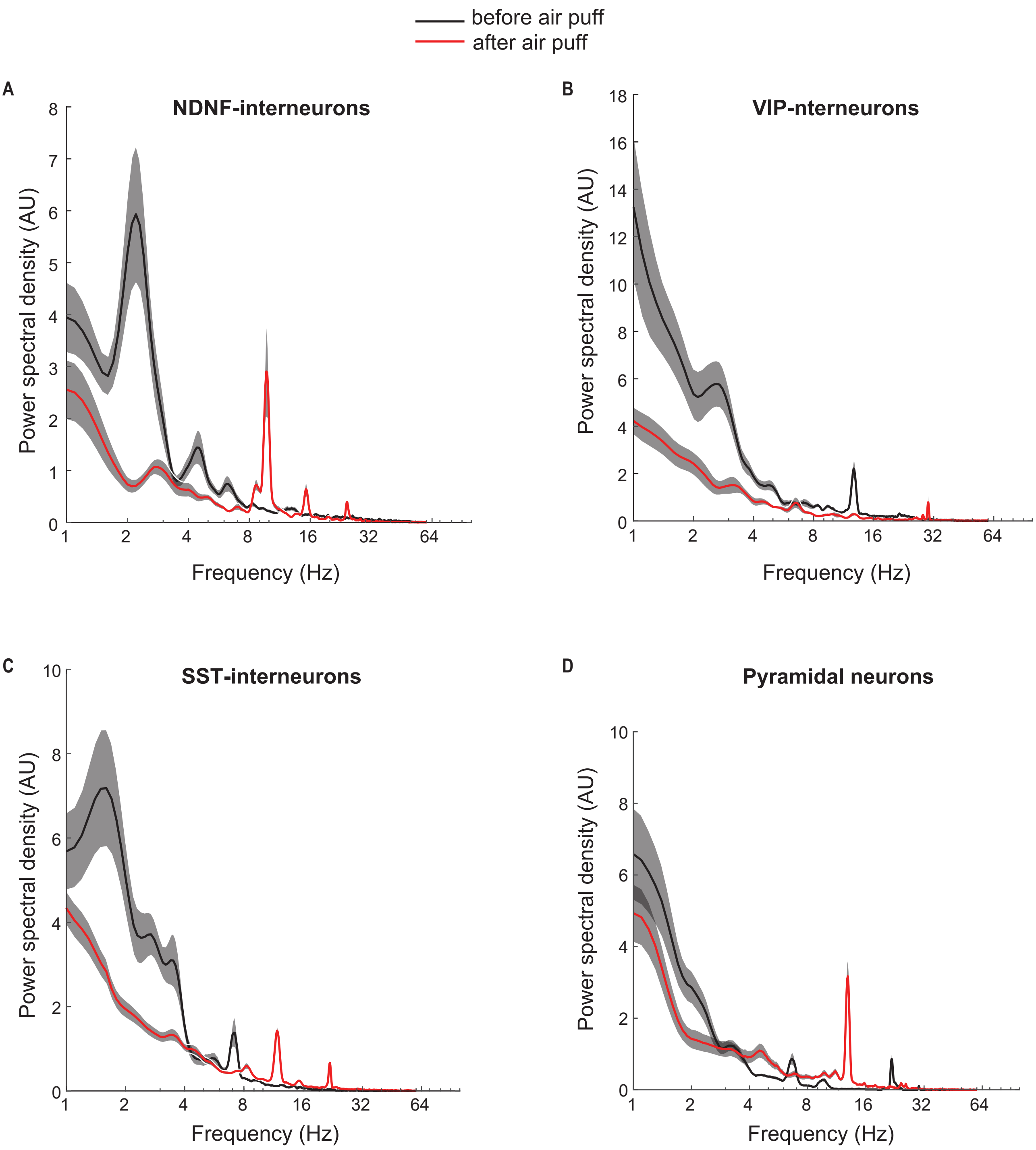


**Figure S15 |** Subthreshold power for distinct V1 cell classes during state transitions.
(A-D) Subthreshold power was computed across different frequencies from the Δ*F/F* traces obtained using Ace-mNeon2 voltage imaging for (A) NDNF-interneurons (n=57 cells), (B) VIP-interneurons (n=44 cells) (C) SST-interneurons (n=42 cells) and (D) a sparsely labeled subset of pyramidal neurons (n=27 cells) during a 3 s time window before and after air puff delivery. The post-air puff period was associated with a decrease in the power of low-frequency and increase in the power of high-frequency subthreshold activity, most prominent in NDNF- and SST-neurons. Power density curves represent mean ± S.E.M.

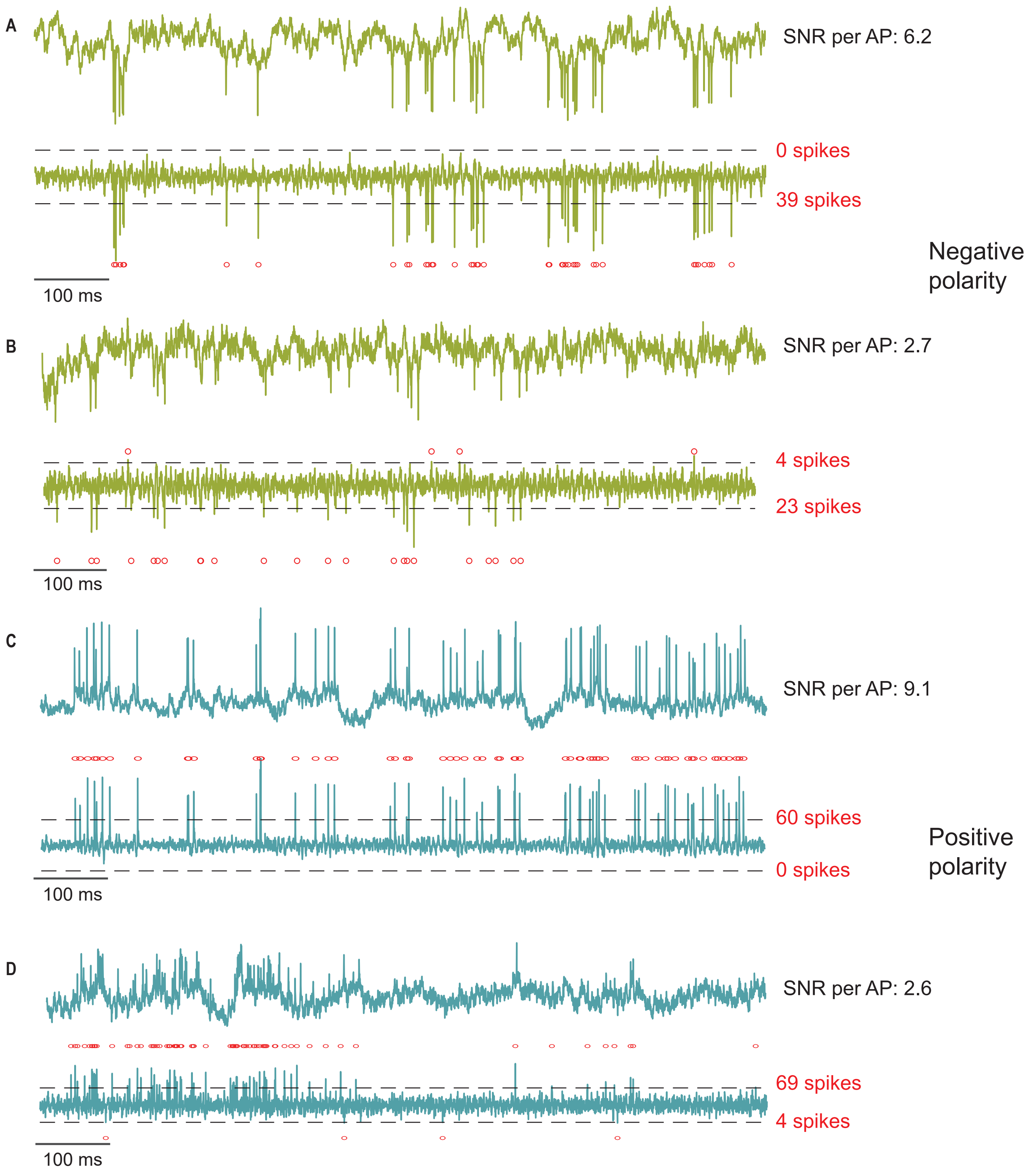


**Figure S16 |** Polarity assignment in V1 DUPLEX recordings in awake mice (see also Methods).
(A) *Top*, representative raw and, *bottom*, spike-identified Δ*F/F* traces from a cell exhibiting negative-polarity spikes at high SNR. Dashed lines indicate 3x S.D. of baseline in either direction. Transients that surpassed this threshold were identified as spikes (red circles) and the direction with the most number of spikes determined the spike polarity. Thus, this neuron was assigned a negative-polarity and the neuron-type was inferred based on Ace-mNeon2 targeting.
(B) Same as (A) for a low SNR trace.
(C) *Top*, representative raw and, *bottom*, spike-identified Δ*F/F* traces from a cell exhibiting positive-polarity spikes at high SNR. Dashed lines indicate 3x S.D. of baseline in either direction. Transients that surpassed this threshold were identified as spikes (red circles) and the direction with the most number of spikes determined the spike polarity. Thus, this neuron was assigned a positive-polarity and the neuron-type was inferred based on pAce targeting.
(D) Same as (C) for a low SNR trace

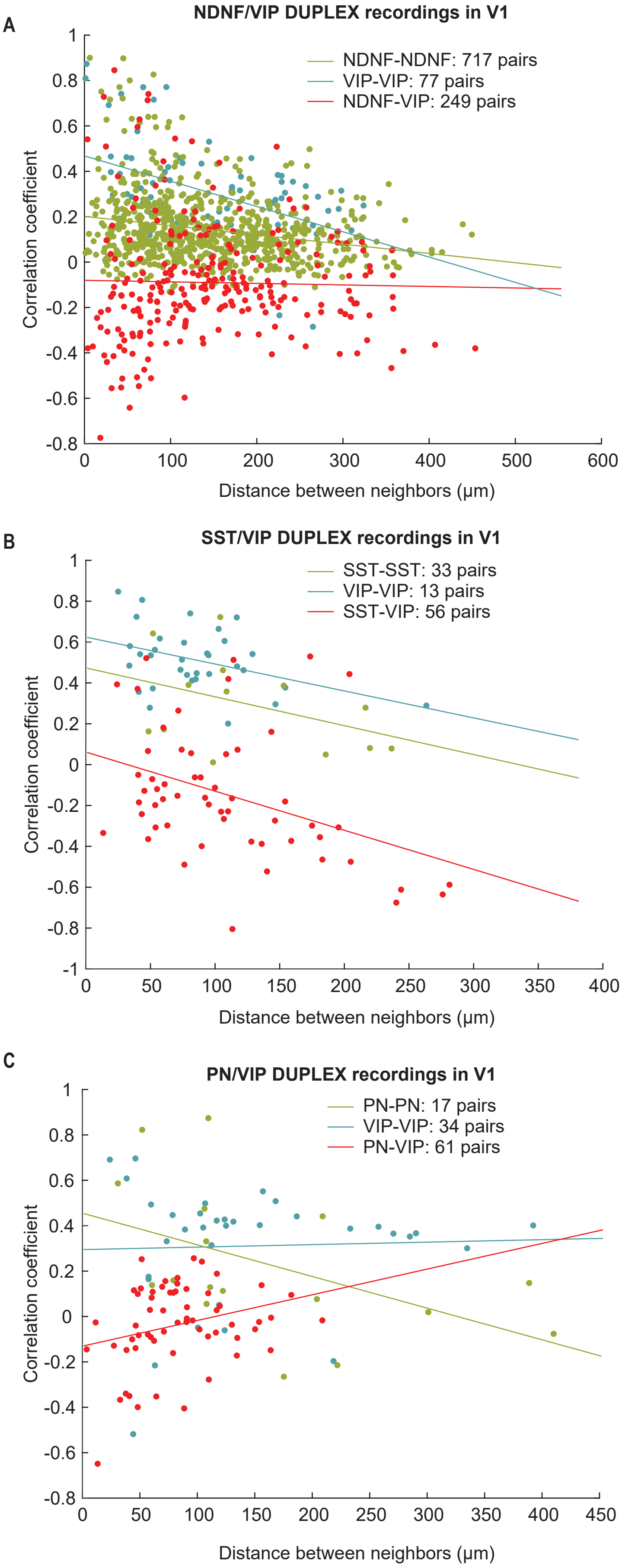


**Figure S17 |** Pairwise correlation coefficients as a function of distance between neighbors in DUPLEX recordings in V1. (A) Zero-time lag correlation coefficients of the $\Delta F/F$ traces between NDNF-NDNF, VIP-VIP and NDNF-VIP cell pairs plotted as a function of distance between the cells in DUPLEX recordings from injected NDNF-Cre+/VIP-Flp+ mice (n=83 NDNF- and 42 VIP-neurons, 5 mice).
(B) Same as (A) for DUPLEX recordings from injected SST-Cre+/VIP-Flp+ mice (n=17 SST- and 25 VIP-neurons, 3 mice).
(C) Same as (A) for DUPLEX recordings from injected AAV-CaMKII-Cre/VIP-Flp+ mice (n=20 PNs and 22 VIP-neurons, 3 mice).

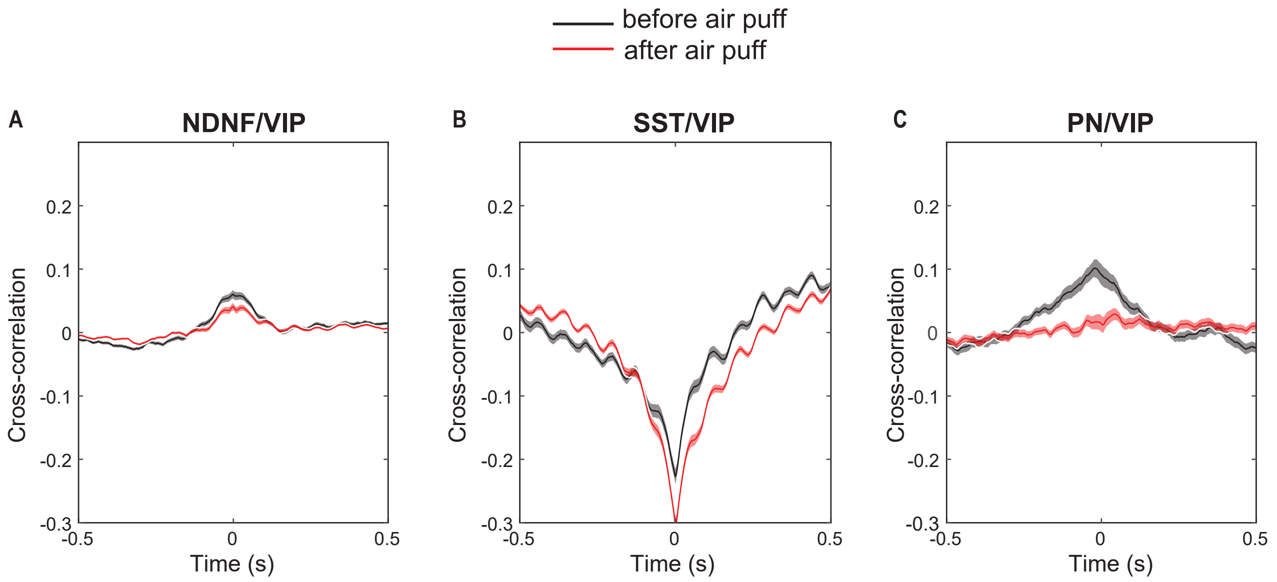


**Figure S18 |** Cross-correlograms of voltage dynamics between distinct V1 cell-types during state transitions. (A-C) Non-zero time lag cross-correlations between the Δ*F/F* traces obtained from DUPLEX recordings before and after air puff in (A) NDNF- and VIP-interneurons (n=351 NDNF-VIP pairs; 31 VIP neurons and 202 NDNF neurons, 5 mice); (B) SST- and VIP-interneurons (n=55 SST-VIP pairs; 23 SST neurons and 16 VIP neurons, 3 mice) and (C) PNs and VIP-interneurons (n=61 PN-VIP pairs; 20 PNs and 22 VIP neurons, 3 mice). Note the small positive correlation between PNs and VIP-cells at negative lag, suggestive of a temporal delay in transmission between the two cell-types. However, no cross-correlation was observed after state change.

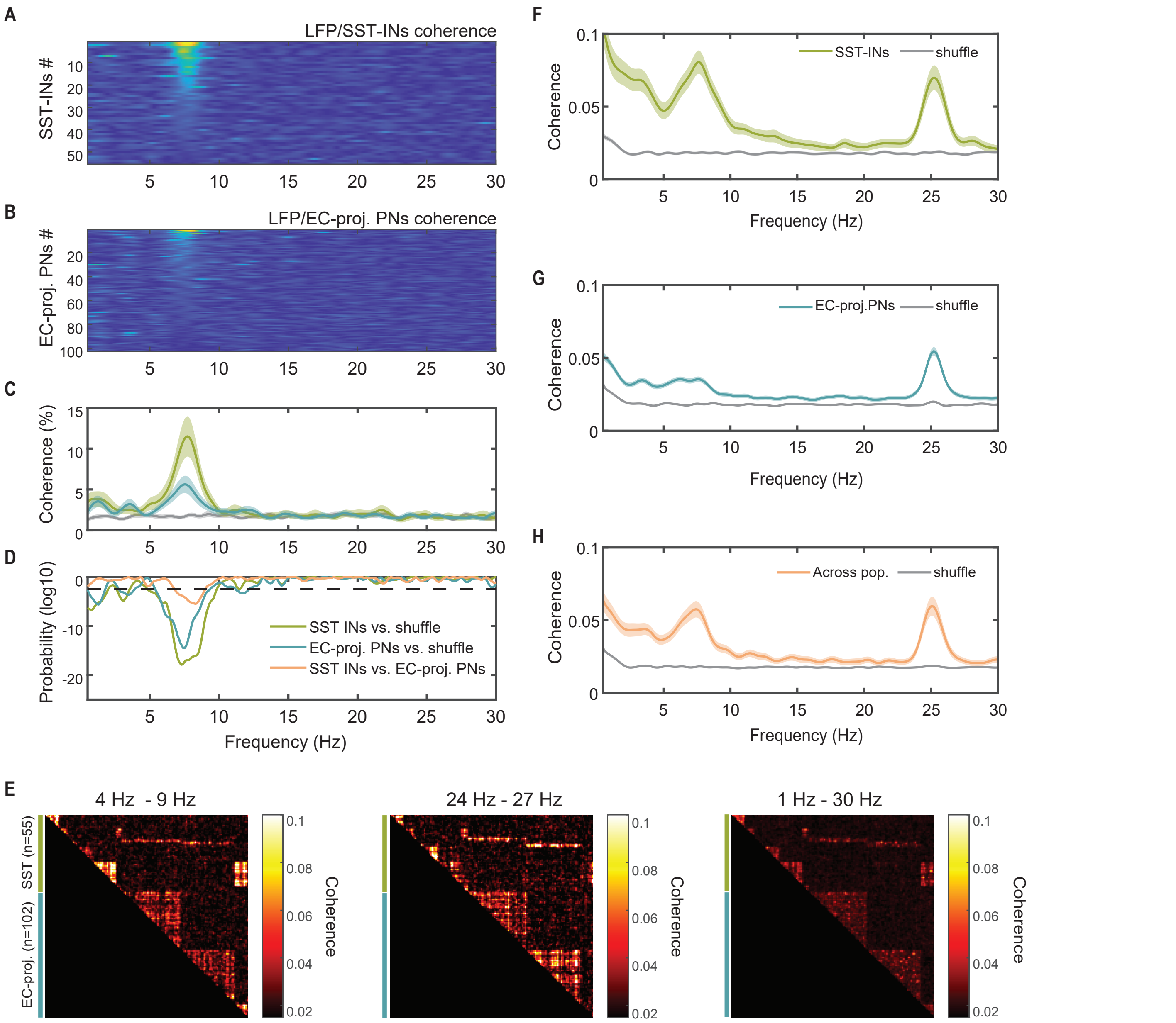


**Figure S19 |** Subthreshold analysis of excitatory versus inhibitory contributions to local field potentials and pairwise coherence of excitatory inhibitory neuronal ensembles.
(A) SST-interneurons (n=55 cells, 6 fields-of-view, 1 mouse) and LFP coherence raster.
(B) EC-projecting PNs (n=102 cells, 6 fields-of-view, 1 mouse) and LFP coherence raster.
(C) Average LFP coherence for each neuronal ensemble.
(D) Probability plot with dashed line corresponding to Bonferroni-corrected significance threshold (p-value<0.01).
(E) Pairwise coherence matrices measured in narrow band theta (4-9 Hz, *left*) and beta (24-27 Hz, *center*) frequencies, as well as in broad band frequency (1-30 Hz, *right*).
(F-H) Average pairwise coherence (F) within SST-neuronal ensembles, (G) within EC-projecting PN ensembles, and (H) between SST-interneurons and EC-projecting PNs.

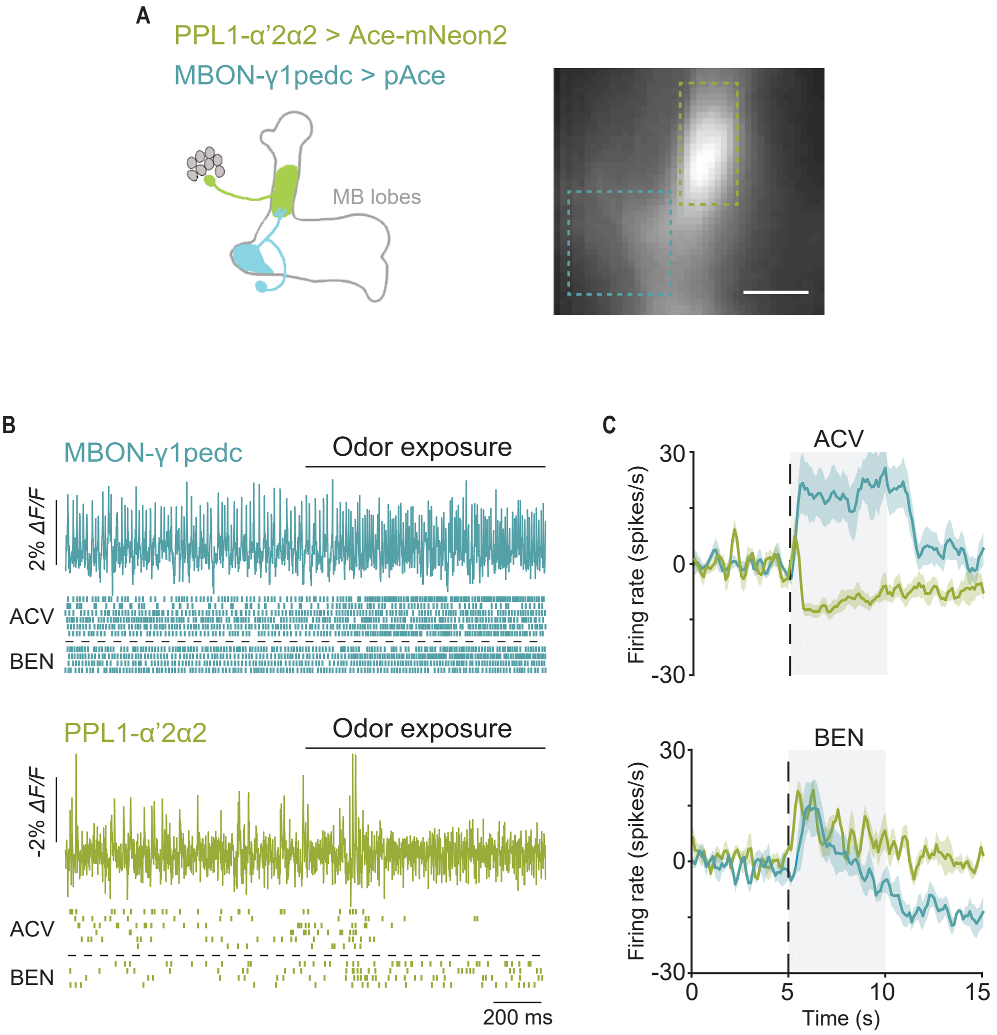


**Figure S20** | Dual-polarity voltage imaging in flies.
(A) *Left*, Cartoon and *right*, epifluorescence image showing expression of Ace-mNeon2 and pAce in two synaptically connected neuron-types for simultaneous dual-polarity voltage imaging. Dashed boxes indicate regions-of-interest comprising the axonal region of PPL1-α’2α2 neuron (green, Ace-mNeon2) and the dendritic region of MBON-γ1pedc >α/β neuron (blue, pAce).
(B) Example traces of evoked spiking during a 2 s time-window around odor onset from a 15-s imaging trial. Raster plots showing responses during 6 trials of exposure to the attractive odorant apple cider vinegar (ACV) and 4 trials of exposure to the repulsive odorant benzaldehyde (BEN) obtained from PPL1-α’2α2 and MBON-γ1pedc>α/β neurons.
(C) Mean spike rate change during ACV (*top*) and BEN (*bottom*) exposure in in the two neurons. (n = 6 trials for ACV; n = 4 trials for BEN; 2 trials per fly).

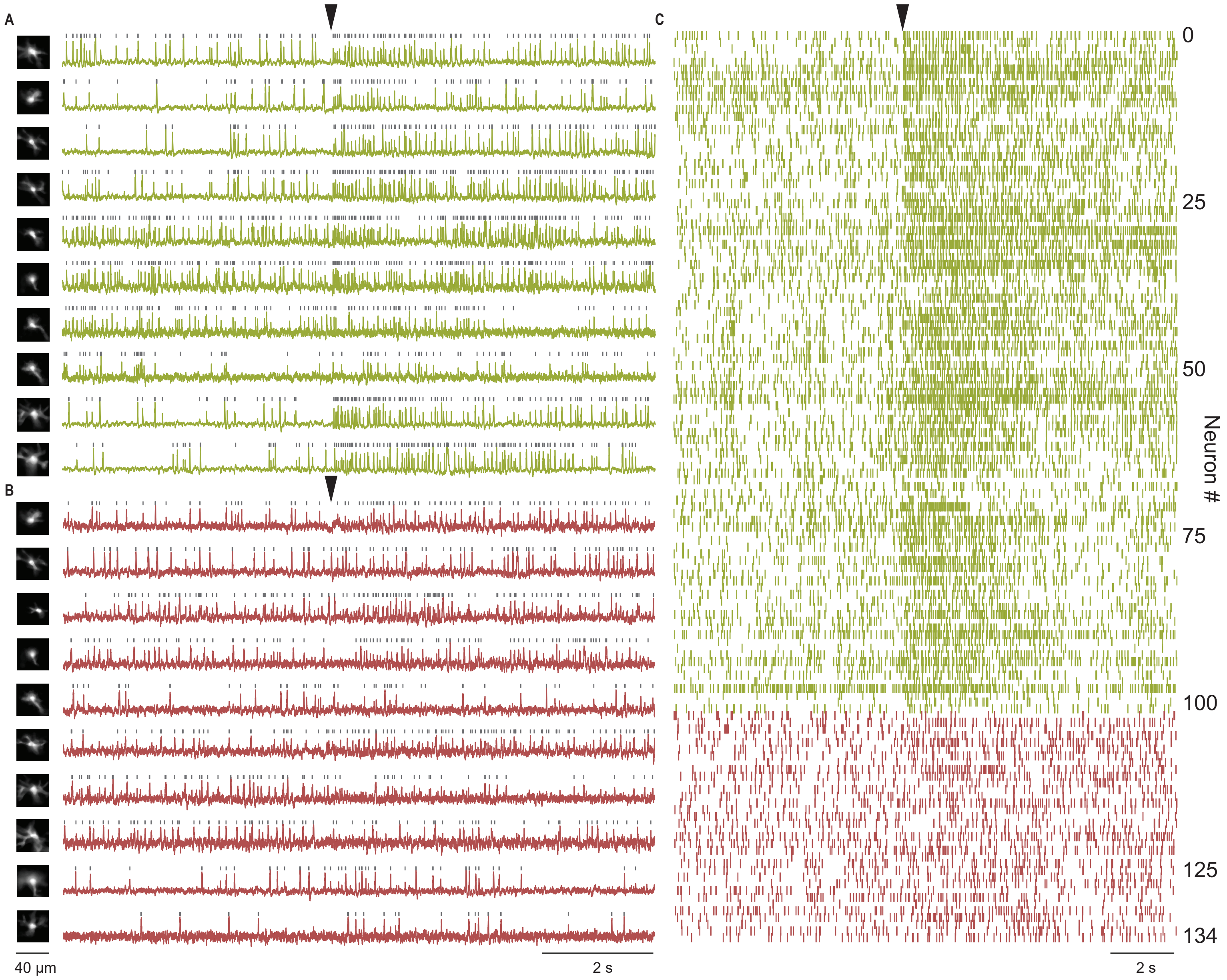


**Figure S21** | Voltage recordings from ACC- and EC-projecting excitatory neurons in dorsal CA1 in an awake behaving mouse during behavioral state transition.
(A) Example Δ*F/F* traces from ACC-projecting PNs retrogradely labeled with Ace-mNeon2.
(B) Example Δ*F/F* traces from EC-projecting PNs retrogradely labelled with VARNAM2.
(C) Summary raster plot for all 134 projection neurons (n=5 fields-of-view, 1 mouse). Arrows indicate rest-to-run transition onset.

| | Activation | | | Deactivation | | |
|---|---|---|---|---|---|---|
| **GEVI** | **$\tau_{fast}$ (ms)** | **$\tau_{fast}$ (ms)** | **% fast** | **$\tau_{fast}$ (ms)** | **$\tau_{fast}$ (ms)** | **% fast** |
| Ace-mNeon2 | 0.77 ± 0.08 | 3.1 ± 0.4 | 67 ± 4 | 0.81 ± 0.03 | 2.8 ± 0.1 | 65 ± 8 |
| VARNAM2 | 0.5 ± 0.19 | 1.9 ± 1.0 | 68 ± 4 | 0.48 ± 0.3 | 3.1 ± 1.8 | 72 ± 7 |
| pAce | 0.51 ± 0.02 | 1.5 ± 0.3 | 82 ± 5 | 0.61 ± 0.03 | 3.2 ± 0.4 | 78 ± 5 |
| pAceR | 2.4 ± 0.3 | 5.4 ± 4.2 | 54 ± 8 | 1.9 ± 1.6 | 7.2 ± 6.8 | 58 ± 4 |

**Table S1 |** Response kinetics of the FRET-opsin indicators.
Responses to depolarizing voltage steps obtained from transfected HEK cells were fitted using a biexponential step function to determine rise and decay kinetics. τfast and τslow indicate time constants of the fast and slow components, respectively. Imaging conditions: 505 nm LED for Ace-mNeon2 and pAce and 565 nm LED for VARNAM2 and pAceR, 25 mW $mm^{-2}$ at sample plane; image acquisition: 3-5 kHz. n=6 cells for Ace-mNeon2, 7 cells for pAce, 4 cells each for VARNAM2 and pAceR. Values represent mean ± S.E.M.
